# Differential Expression of α, β, and γ Protocadherin Isoforms During Differentiation, Aging, and Cancer

**DOI:** 10.1101/2021.03.07.434314

**Authors:** Michael D. West, Ivan Labat, Jie Li, Pam Sim, Jeffrey Janus, Hayley Mangelson, Shawn Sullivan, Ivan Liachko, Paul Labhart, Maddy Craske, Brian Egan, Karen B. Chapman, Nafees N. Malik, Dana Larocca, Hal Sternberg

## Abstract

The cadherin family of cell surface glycoproteins plays a fundamental role in cell-cell recognition, thereby participating in diverse biological process such as embryonic morphogenesis and oncogenic transformation. The subset of clustered protocadherin (PCDH) genes generated from the α, β, and γ loci, have been widely studied for their potential role in neuronal cell-cell recognition and neurogenesis, however their broader role in normal embryonic development and cancer has not been examined in detail. We utilized human embryonic stem (hES) cells to model early human development *in vitro*, comparing PCDH isoform transcription in diverse types of embryonic progenitors with normal adult-derived and cancer counterparts. Embryonic progenitors express genes from the α and β cluster at levels comparable to that seen in the CNS, while fetal and adult-derived cells express primarily from the γ cluster. Replicative senescence left fibroblasts with markedly lower expression of all isoforms. We observe that an embryonic pattern of clustered protocadherin gene expression and associated CpG island methylation is commonly associated with cancer cell lines from diverse tissue types. The differential regulation of the α, β, and γ loci coincide with alternate regions of DNA accessibility at CTCF binding sites and lamina-associated domains and CPL expression correlated with the expression of *LMNA* and *LMNB1*. These observations support a potential role for the differential regulation of genes within the clustered protocadherin locus in selective cell-cell adhesion during embryogenesis, regeneration, cancer and aging.

## Introduction

Early research into the cellular basis of embryology suggested that the complexities of tissue morphogenesis must employ precise mechanisms for cell-cell recognition and differential adhesion^1, 2^. Such a program seems necessary for the generation of a uniformity of cell types, association with specific neighboring cells, and the formation of discrete boundaries. Evidence for such cell-cell recognition and adhesion began with the pioneering studies of H.V. Wilson who first reported the spontaneous reassociation and regeneration of sponges from dissociated cells^3^.

The evolution of more complex organisms carrying an increasingly broad array of differentiated cell types arguably would depend to an even greater extent on this phenomenon. Johannes (Hans) Holtfreter referred to this selective adhesion as “tissue affinity” and in 1939 theorized that, “… attraction and repulsion phenomenon is operating between various cell types during development and that information on this system will yield valuable information concerning the shiftings and segregations of tissues during organogenesis.” As a result of such early studies, the elucidation of the role of differential cell-cell adhesion in morphogenetic process became widely recognized as a milestone in developmental biology comparable to the discovery of organizers by Spemann and Mangold^4^.

In the 1950s, experimental embryologists extended these studies to chick embryonic development. The regenerative nature of embryonic anlagen was widely utilized in experimental embryo transplant studies, such as the cross-transplantation of embryonic tissues from quail to chicken. Studies of disaggregated limb bud chondrogenic and myogenic cells as well as embryonic mesonephros and retina could re-associate and regenerate scarlessly, however, aggregation markedly decreased later in the course of development^5, 6^. This led to the question of the “why” and “how” of the developmental repression of cell-cell adhesion regulation during embryogenesis and the loss of regenerative potential.

In an attempt to answer the question as to “why” natural selection led to the repression of regeneration during development followed by aging, George Williams suggested the model of “antagonistic pleiotropy”, in part as an explanation of why “after a seemingly miraculous feat of morphogenesis a complex metazoan should be unable to perform the much simpler task of merely maintaining what is already formed^7^.” The theory posits that some traits selected for based on their value early in the reproductive period of the life cycle, have adverse effects in the post-reproductive years. One example is the nearly global repression of telomerase expression early in embryonic development and the re-expression of the gene *(TERT)* in ∼90% of cancer types^8^. The repression of *TERT* may reduce the risk of malignancy early in life, but may lead to cell senescence late in life.

The “how” of the regulation of differential embryonic cell adhesion was first envisioned by William Dreyer, William Gray, and Leroy Hood in 1967 in a largely theoretical paper on the molecular basis of immunoglobulin diversity. They proposed a complex family of cell adhesion molecules may have been co-opted during evolution to create diverse antibodies.^9^” In 1999, William Dreyer proposed that retrotransposition events evolved a complex family of “area code” cell adhesion molecules that potentially include immunoglobulins, protocadherins, and olfactory receptors that regulate the complexities of development^10^.

The characterization of the clustered protocadherin locus (CPL), revealed such a predicted complex family of contiguous genes that like the immunoglobulin loci, contains numerous variable exons that can combine with constant exons to create a vastly complex array of cell adhesion interfaces^11^. The superfamily member genes are arranged in three clusters designated Pcdhα, Pcdhβ, and Pcdhγ. The α and γ clusters possess variable exons encoding cadherin ectodomains, one transmembrane, and short cytoplasmic domains. In contrast, the β cluster contains unique single exon genes. In *Homo sapiens* there are 15 α, 15 β, and 22 γ domains. Each variable domain is preceded by a promoter region. These proteins generate homophilic associations in trans as evidenced by the induction of cell aggregation when PCDH constructs are expressed in K562 cells^12, 13^. In neurons, a vast complexity can be generated by stochastic activation of isoforms in a biallelic fashion^14^. Cis-combinations of the PCDH γ genes alone have been estimated to generate a potential diversity of >10^5^ diversity in unique cell interfaces.

Following up on his hypothesis, William Dreyer proposed in 1999 that retrotransposition events evolved a complex family of adhesion molecules that regulate the development of the CNS, including the synapses in the olfactory bulb. Subsequently, the majority of research on CPL homophilic interactions has focused on neuronal self-recognition^15^ and circuit assembly^12^. While the role of cadherins in cell adhesion and development is well-established^16^, the potential role of the CPL in morphogenesis and epimorphic regeneration outside of the CNS has not been elucidated.

The isolation of human embryonic stem cell lines provides an *in vitro* source of diverse human cell types including those capable of embryonic cell adhesion as evidenced by their spontaneous formation of embryoid bodies and organoids^17, 18^. Utilizing clonally-purified progenitor cell lines as a model the early regenerative phenotype of cells before the embryonic-fetal transition (EFT), we applied deep neural network algorithms to transcriptomic data from relatively large numbers of embryonic vs adult cell and tissue samples and reported putative genes differentially expressed before and after the EFT^19^. Among the pre-EFT genes was the cell adhesion gene, *PCDHB2*. Here we compare gene expression in diverse *in vitro* embryonic progenitors compared to adult normal and cancer cell lines and report a striking differential expression of CPL isoforms in this *in vitro* model of human embryonic development. In addition, we observe a shift in CPL gene expression in diverse cancer cell types toward embryonic patterns of expression consistent with that predicted by the theory of antagonistic pleiotropy.

## Results

Pluripotent stem cell (PSC)-derived progenitors such as those derived clonally, display markers of primitive embryonic anlagen despite extensive passing or differentiation *in vitro*^19, 20^. We therefore designated the lines “clonal embryonic progenitor cells” to distinguish them from fetal and adult cell counterparts. These cell lines represent primarily diverse stromal and parenchymal progenitors, as opposed to epithelial cell types, therefore we will designate these diverse clonal embryonic progenitor cell lines as “EPs” as compared to stromal fetal cells (“FCs”), and stromal adult non-epithelial lines (“ANE”). We utilized EPs and their fetal and adult cell type counterparts as an *in vitro* model of cells displaying a pre- and post EFT phenotype.

### Clustered Protocadherin Isoforms are Differentially Expressed in Embryonic vs Fetal and Adult Non-Neuronal Somatic Cell Types

RNA-sequence data was obtained from diverse human embryonic, adult, and cancer cell types, specifically including: four different human ES cell lines and two iPSC lines (“PC” cells); 42 diverse clonal EP cell lines; eight FCs including three brown preadipocyte cultures and five fetal skin dermal fibroblasts spanning 8-16 weeks of development; 89 diverse stromal and parenchymal non-epithelial cell types (ANE), and five adult neuronal cell (NC) types including neurons and astrocytes (see Supplementary Table I for cell type descriptions). Since EP cells display a pre-fetal pattern of gene expression, we refer to them as “embryonic” contrasted with that of pluripotent stem cells (PCs) and data relating to diverse ANEs and adult epithelial cells (AECs) to which we refer as “adult.”

To identify global developmental alterations in gene expression that encompass numerous differentiated cell types that may account for the pre-EFT regenerative phenotype, we filtered the isoform data for highly statistically-significant gene expression differences distinguishing embryonic from adult cells regardless of differentiated cell type. As shown in Figure 1, 1079 isoforms showed a significant difference in expression (adjusted p value <0.05) in the EP vs ANE cells, with many of the most significant and largest fold-changes being isoforms of the CPL (Figure 1) Embryonic up-regulated CPL isoforms were typically from the α and β clusters, while adult up-regulated isoforms were from the γ cluster.

**Figure 1.**
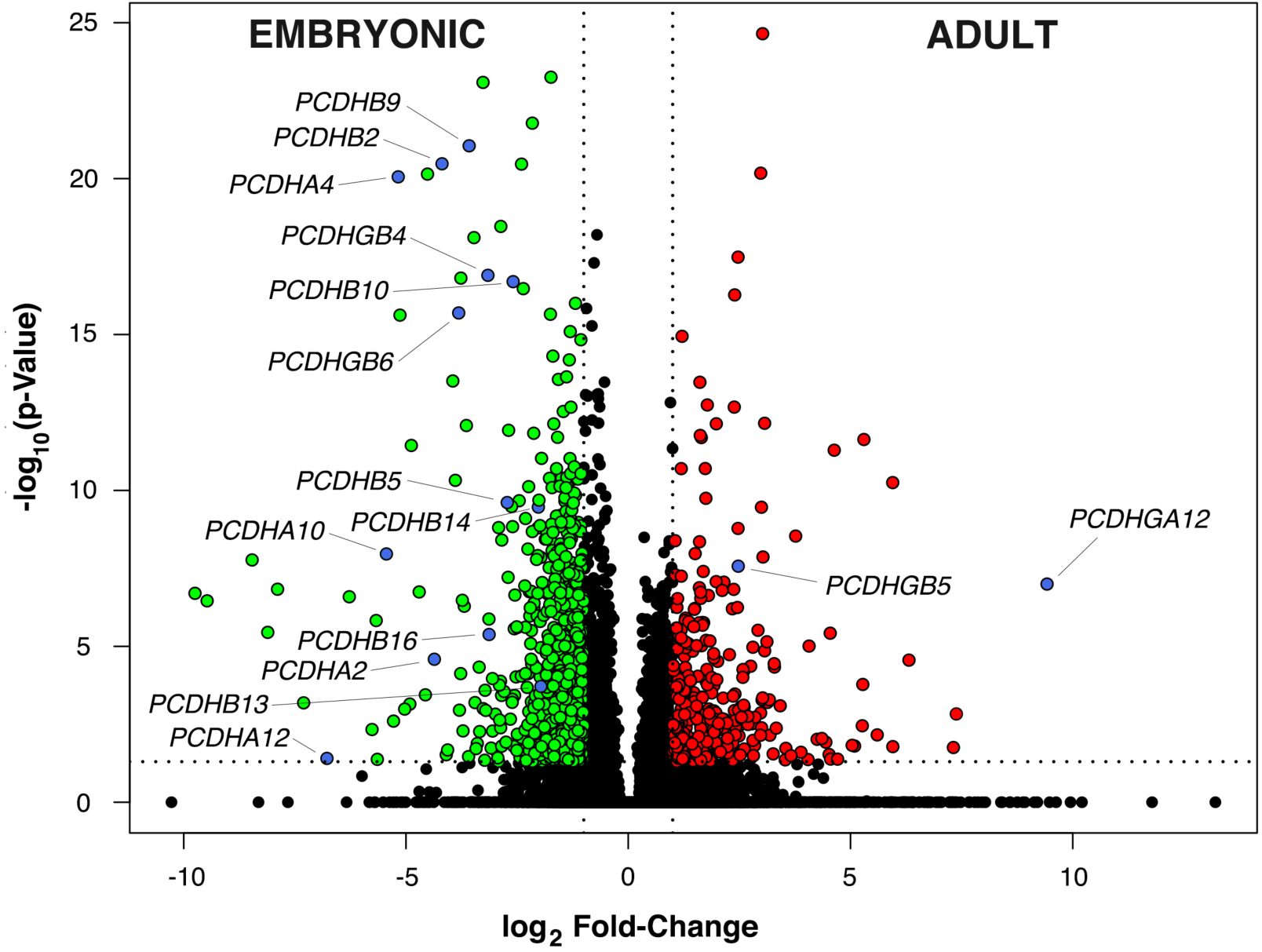
Volcano plot of differentially-expressed transcript isoforms determined by RNA-seq. Log_2_ fold-change (FC) versus -log_10_ (Bonferroni-adjusted p-value) is plotted. The underlying data is derived from RNA sequencing FPKM values for 42 diverse human clonal embryonic progenitor cell lines and 92 diverse adult-derived stromal and parenchymal cell types. The horizontal dotted line indicates a linear x-adjusted p-value of 0.05 and vertical dotted lines indicate a linear fold-change (FC) of 2. Transcripts passing the cutoff criteria of p < 0.05 and FC > 2 (elevated expression in adult) are represented with red points; transcripts with p < 0.05 and FC < -2 (elevated expression in embryonic) are represented with green points. Points labelled in blue are CPL isoforms.

We chose for further study a representative member from each of the α and β loci (*PCDHA4* and *PCDHB2)* that were significantly up-regulated in the majority of embryonic cells compared to adult counterparts (p-values of 8.8 x 10^-21^ and 3.3 x 10^-21^ respectively), and showed mean fold-changes of embryonic versus adult expression (FPKM) of 36.1 and 18.2 respectively (Figures 2a-c). From the γ locus, the isoform *PCDHGA12* showed adult up-regulation in the majority of adult cell types with an adjusted p-value of 1.0 x 10^-7^ and a mean fold-change in adult versus embryonic expression (FPKM) of 686-fold. While isoform expression from the α and β clusters appeared to be often up-regulated in embryonic (PCs and EPs) compared to adult (ANEs), isoforms from the γ locus were heterogeneous with *PCDHGB4* and *PCDHGB6* being up-regulated in embryonic cells while *PCDHGB5* and *PCDHGA12* being up-regulated in adult counterparts (Figure 2).

**Figure 2.**
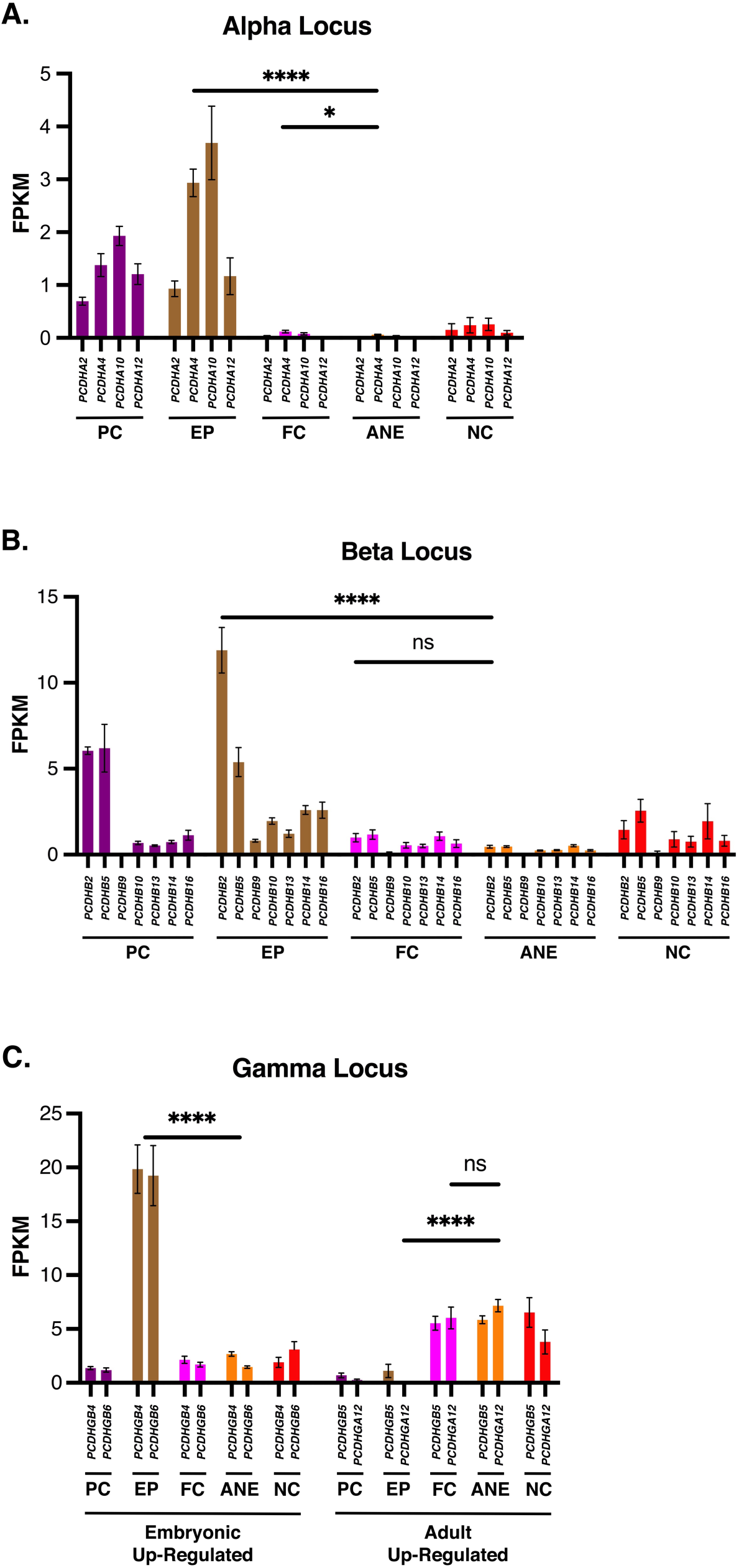
Differential expression of CPL genes in embryonic vs adult cell types. A). Mean expression in FPKM of RNA-seq reads of *PCDHA2, PCDHA4, PCDHA10,* and *PCDHA12* in diverse pluripotent stem cell (PC), hES-derived clonal embryonic progenitors (EP), fetal-derived cells (FC), adult-derived cells (AC), and neuronal cell (NC) types. B). Mean expression of *PCDHB2, PCDHB5, PCDHB9, PCDHB10, PCDHB13, PCDHB14, PCDHB16* in aforementioned cell types. C) Mean expression of *PCDHGB4, PCDHGB5, PCDHGB6,* and *PCDHGA12* in diverse embryonic, adult, and cancer cell types. ES and iPS cell lines include four different human ES cell lines and two iPS cell lines; Diverse EPs include 42 diverse human clonal embryonic progenitor cell lines, fetal-derived cells include three cultured of brown preadipocytes and five fetal arm skin fibroblast cultures from 8-16 weeks of gestation; Diverse Normal Somatic Cells include 87 diverse stromal and parenchymal cell types; five neuronal cell cultures including neurons and diverse astrocyte cell types; Epithelial cells include 25 diverse human epithelial cell types; Sarcomas include 39 diverse human sarcoma cell lines; Carcinomas include 33 diverse carcinoma and adenocarcinoma cell lines; (See Supplementary Table I for complete cell type descriptions.) (ns – not significant) (* p<0.05) (** p<0.01) (*** p<0.001) (**** p< 0.0001). Error bars S.E.M.

### The Onset of Adult CPL Isoform Expression Occurs at or Before the Embryonic-Fetal Transition

The EFT is commonly associated with a loss of the capacity for scarless regeneration in numerous tissues of the mammalian body. In the case of human skin, this appears to coincide approximately with Carnegie stage 23 (eight weeks of gestation). We therefore tested the hypothesis that the transition from an embryonic scarless regenerative phenotype to a fetal/adult non-regenerative state correlates temporally with altered CPL isoform expression. We examined the expression of CPL isoforms in early passage fibroblasts from the medial aspect of the upper arm synchronized in quiescence (Fetal Cells (FCs)) in comparison with EP and ANE counterparts (Figure 2). In the case of the representative genes *PCDHA4*, *PCDHB2*, and *PCDHGA12*, CPL isoform expression in 8-16 week-old FCs appeared exhibited an adult-like pattern of expression with the possible exception of *PCDHA4* where FCs showed a modestly-significant elevation over adult counterparts (ANE cells) (p< 0.05). A broader analysis of 34 early passage dermal fibroblasts from adults aged 11-83 years all showed an adult pattern (low *PCDHA4*, low *PCDHB2*, and high *PCDHGA12* with no significant alteration with during aging *in vivo* (data not shown). We conclude that the transition in CPL expression may therefore occur before the EFT (Human Carnegie stage 23) *in vivo*.

PC cells such as embryonic stem (ES) and induced pluripotent stem (iPS) appeared to display a subset of isoforms including *PCDHA2, PCDHA4, PCDHA10, PCDHA12, PCDHB2, PCDHB5*, and like EPs, express low levels of the adult markers *PCDHGB5* and *PCDHGA12* (Figure 2). Importantly, the expression level of the CPL isoforms up-regulated in embryonic cells (both PCs and EPs) was comparable to the CNS-derived cell types used in this study which included neurons and astrocytes of diverse origin. Given the evidence in support of the role of CPL isoforms in the development of the CNS, the comparable or even greater levels of transcripts for CPL isoforms in diverse somatic cell types outside the CNS in EPs is consistent with a hypothesis that CPL isoforms are expressed in the embryonic (pre-EFT) state, and may therefore play a role in cell-cell recognition during development.

### Diverse Cancer Cell Lines Commonly Display an Embryonic Pattern of CPL Isoform Expression (Embryo-Onco Phenotype)

Shared gene expression changes that occur in diverse somatic cell types after completing embryonic organogenesis may reflect antagonistic pleiotropy. That is, the post-EFT repression of pathways critical for cell-cell recognition and morphogenesis may limit regeneration but have the selective advantage of providing effective tumor suppression. We previously demonstrated such a role for the catalytic component of telomerase *(TERT)* by screening for evidence of repression across diverse normal somatic cell types together with their malignant counterparts^8^. We therefore performed a similar survey using RNA-seq-based transcriptomic data of the previously-described PCs, diverse EPs, diverse ANE cell types as well as 24 diverse normal human adult epithelial cell (AEC) types, 39 diverse human sarcoma cell (SC) lines, and 35 diverse carcinoma and adenocarcinoma cell (CAC) lines.

As shown in Figure 3, the representative embryonic CPL isoform transcript markers *PCDHA4* and *PCDHB2* are significantly down-regulated in ANE cells and AECs compared to EPs (p< 0.0001). Similarly, *PCDHGA12* expression was significantly elevated in ANE cells and AECs compared to EPs (p< 0.0001). All isoforms significantly differentially expressed are shown in Supplementary Figures 1A, B, and C. Interestingly, diverse cancer lines such as SCs and CACs (representing cancers originating from stromal and epithelial cell types respectively) when compared to normal ANE and AEC cultures respectively showed a highly significant shift toward an embryonic pattern of CPL gene expression. In the case of the α and β loci, *PCDHA4* and *PCDHB2* are significantly up-regulated in SCs and CACs compared to normal ANE and AEC counterparts (p< 0.01 and p< 0.05 respectively in the case of *PCDHA4* and p< 0.0001 and p< 0.001 respectively in the case of *PCDHB2*). Similarly, *PCDHGA12* expression was significantly reduced showing a more embryonic pattern in SCs and CACs (p<0.001 for SCs compared to normal ANE counterparts and p<0.05 for CACs compared to normal AEC counterparts).

**Figure 3.**
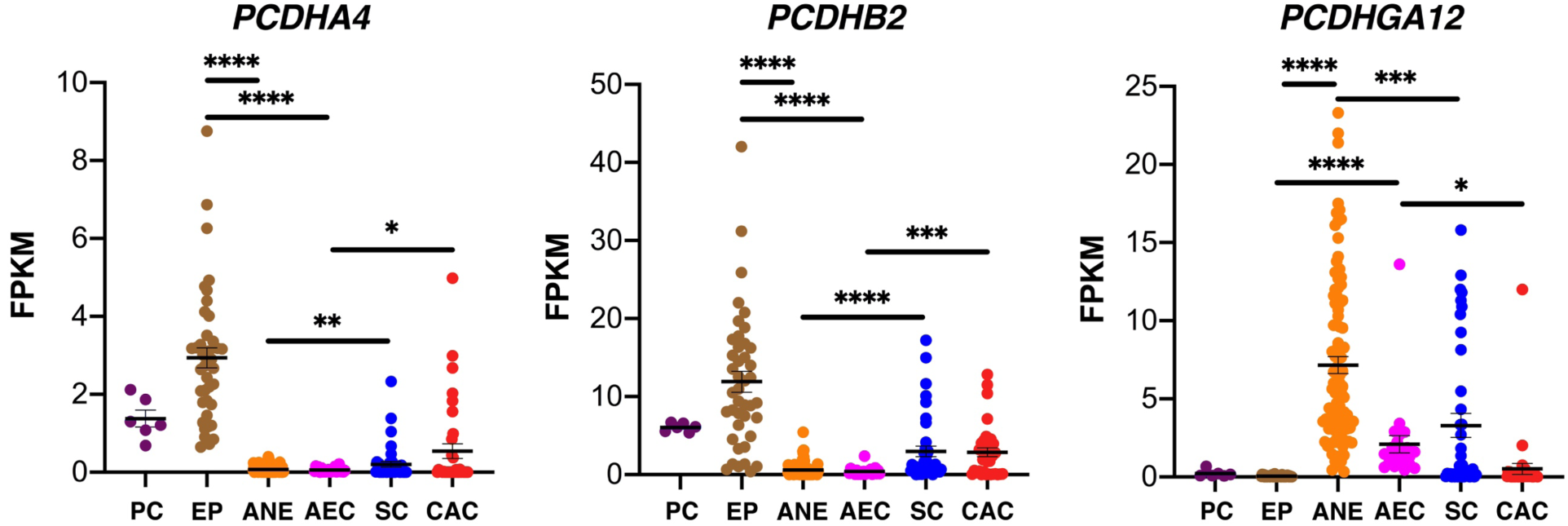
Differential expression of *PCDHA4, PCDHB2,* and *PCDHGA12* in diverse embryonic, adult, and cancer cell types. Mean expression in FPKM of RNA-seq reads from pluripotent stem cell lines include four different human ES cell lines and two iPS cell lines (PC); 42 diverse hES-derived clonal embryonic progenitors (EP); Adult non-epithelial (ANE) cells include 97 diverse stromal and parenchymal cell types; Adult epithelial cells (AEC) include 25 diverse human epithelial cell types; Sarcoma cell (SC) lines include 39 diverse sarcoma cell types; Carcinomas and adenocarcinoms (CAC) include 33 diverse carcinoma and adenocarcinoma cell lines. (See Supplementary Table I for complete cell type descriptions.) (ns – not significant) (* p<0.05) (** p<0.01) (*** p<0.001) (**** p< 0.0001). Error bars S.E.M.

While both sarcoma and carcinoma cell lines appear to frequently display an embryonic pattern of CPL gene expression (i.e. up-regulated *PCDHA4* and *PCDHB2* and down-regulated *PCDHGA12*), the combination of individual isoform expression in the lines appear heterogeneous. This is consistent with the hypothesis that CPL isoforms play a dual role in both differentiated cell type-specific cell-cell recognition as well as regulating alterations unique to the EFT. In addition, sarcoma cell lines appeared heterogeneous in regard to their embryonic vs adult-like CPL isoform profile, carcinoma more frequently embryonic. In the case of the marker *PCDHGA12*, the majority of normal adult cells expressed the isoform while the majority of cancer cell lines did not (Supplementary Figure 1C). For example, using the cutoff of 0.5 FPKM as a lower limit of expression, only 2/97 adult-derived stromal and parenchymal cell types (hepatocytes in both cases), and 1/23 cultured epithelial cell types did not express *PCDHGA12*. However, 21/39 (54%) of sarcoma lines and 33/35 (94%) of carcinoma and adenocarcinoma cell lines did not express *PCDHGA12* at a level of at least 0.5 FPKM. In a manner unique to cancer cell lines, 6/39 (15%) of sarcoma and 5/35 (14%) of carcinoma lines expressed *PCDHA1* at a level > 0.5 FPKM, while no PCs, EPs, FCs, ANEs, or BCs expressed the gene at that threshold level (data not shown).

To determine whether the embryo-onco phenotype extends beyond the CPL, we examined the correlation of *PCDHA4, PCDHB2,* and *PCDHGA12* expression with *COX7A1*, a robust marker of the EFT in many stromal and parenchymal cell types^19^. *COX7A1* expression commences at approximately the EFT, plateauing in adulthood, and is not expressed in the majority of cancer cell lines^19^. As shown in Figure 4, the adult marker *COX7A1* appears to be inversely correlated with the embryonic markers *PCDHA4* and *PCDHB2* while showing a trend toward directly correlating with the expression of the adult marker *PCDHGA12* in diverse sarcoma cell lines (R^2^ = 0.52, p<0.0001). These data therefore suggest that the majority of sarcoma, carcinoma, and adenocarcinoma cell lines display an embryonic pattern of CPL isoform expression, correlating with a down-regulation of *COX7A1* in support of an embryo-onco phenotype inclusive of the altered expression of multiple genes.

**Figure 4.**
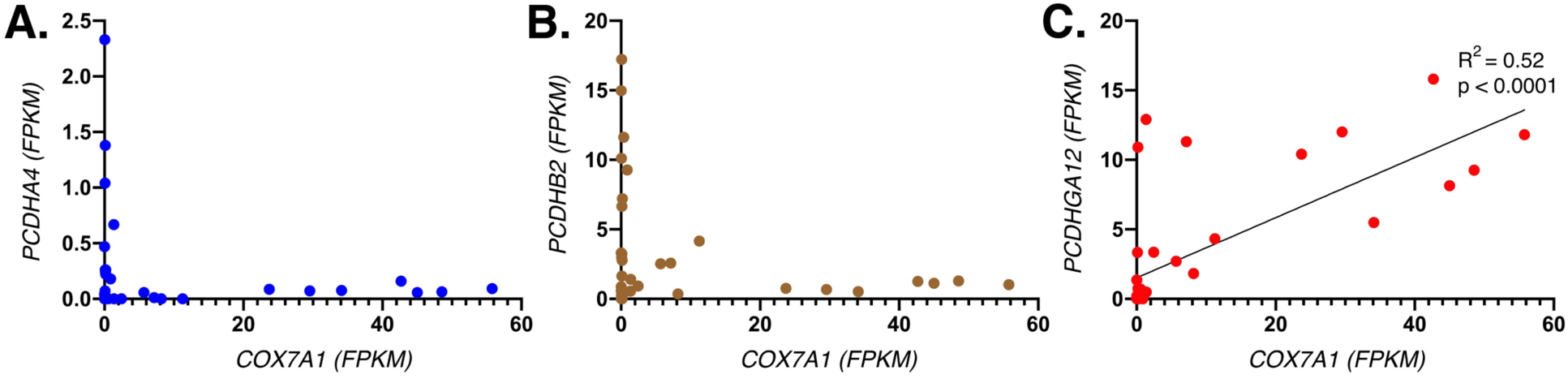
Correlations of embryonic and adult markers in sarcoma cell lines. FPKM values of the embryonic markers A) *PCDHA4,* B) *PCDHB2* and C) *PCDHGA12* are plotted against the adult marker *COX7A1* in 39 diverse sarcoma cell lines.

### Altered Embryonic vs Adult Gene Expression Coincides with Modifications in Chromatin Accessibility, CTCF Binding, and Hypermethylated CpG Islands

We next examined the epigenetic status of the α, β, and γ loci beginning with ATAC (Assay of Transposase Accessible Chromatin) sequencing to identify accessible regions of chromatin and potential interactions with DNA binding proteins^21^. Accessibility coincided with expressed CPL genes in embryonic and adult counterparts of osteogenic mesenchyme and vascular endothelium as determined by the mRNA read coverage (Figure 5). ATAC accessibility also frequently coincided with CTCF footprints as determined by TOBIAS (a computational tool using ATAC sequence data allowing determination of genome-wide transcription factor binding dynamics relative to chromatin organization). CTCF has previously been reported to participate in gene regulation in the CPL by regulating the structure of chromatin domains and regulating cis interactions with enhancers^22^.

**Figure 5.**
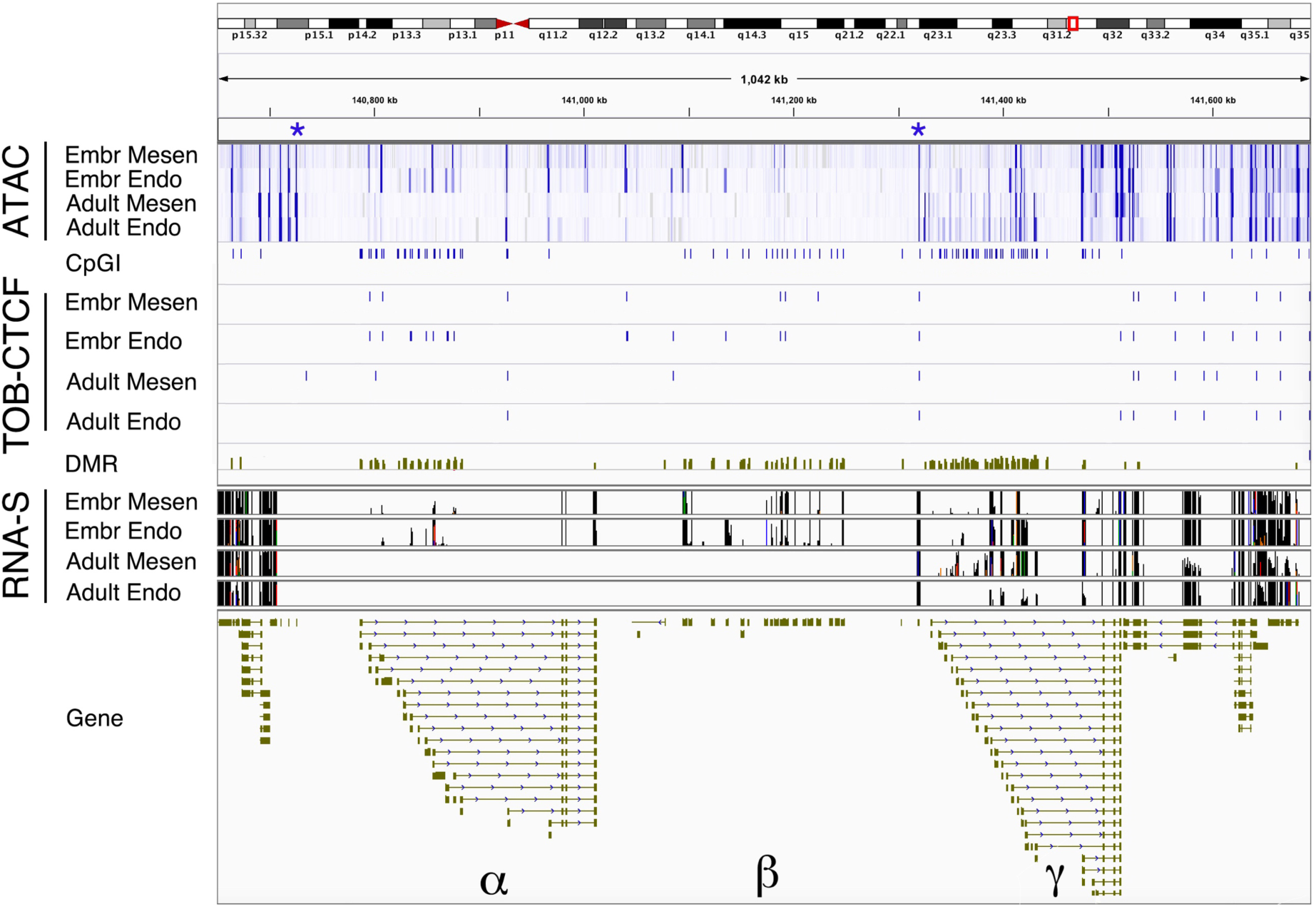
IGV view of chromosomal features and RNA-sequence reads from the CPL. Rows represent: 1) ATAC-seq of the embryonic progenitor osteogenic mesenchymal line 4D20.8 (Embr Mesen); the embryonic vascular endothelial line 30-MV2-6 (Embr Endo); adult-derived osteogenic mesenchyme (MSCs); and adult-derived human aortic endothelial cells (HAECs). Blue asterisks make region of apparent decreased accessibility in adult cells; 2) CpG islands (CpGIs) downloaded from UCSC (cpgIslandExt.hg38.bed) where length > 200 bp and > 60% of expected CpGs based on G and C content; 3) CTCF binding sites in Embr Mesen, Embr Endo, Adult Mesen, and Adult Endo based on TOBIAS analysis of ATAC footprints; 4) Differentially Methylated Regions (DMRs) are shown where elevated green represents hypermethylation in embryonic cells and depressed red signal represents relative hypomethylation in embryonic compared to adult cells; and 5) RNA sequence reads for the alpha, beta, and gamma loci. (hg38 Chromosome position: Chr5: 140,650,000-141,700,000), image captured from IGV.

We next examined CpG methylation in the α, β, and γ loci as determined by Whole Genome Bisulfite Sequencing (WGBS). Four hES cell-derived clonal EP cell lines were sequenced together with their respective adult-derived counterparts, namely: 4D20.8, a clonal embryonic osteochondral progenitor^23^ and adult bone marrow-derived mesenchymal stem cells (MSCs); 30-MV2-6, a clonal embryonic vascular endothelial cell line together with adult-derived aortic endothelial cells (HAECs); SK5 a clonal embryonic skeletal muscle progenitor together with adult skeletal myoblasts; and E3, an embryonic white preadipocyte cell line^24^ together with adult subcutaneous white preadipocytes. Differentially-methylated regions (DMRs) were identified based on significant (p <0.05) difference in methylated CpGs in three of four of the EP/ANE pairs using Metilene software. DMRs in the CPL are shown in Figure 5. DMRs co-localized with the first exon of α and γ isoforms and the gene body of β isoforms, with CpG islands (CpGI), the pattern of read coverage, and surprisingly, were uniformly hypermethylated in embryonic cells compared to adult counterparts regardless of whether the isoform was embryonic or adult-specific. Of the 21,317 significant DMRs identified in the entire genome, the majority (20,022) were hypermethylated (data not shown), however, overall genomic CpG methylation was not significantly different in EPs vs their adult counterparts (Supplementary Figure 2).

### Embryonic DMRs within the CPL are Observed in Diverse Cancer Cell Lines and Appear Distinct from CIMP

We next determined DMR methylation status in sarcoma cell (SC) lines from cell types corresponding to the EPs and ACs of mesenchymal, endothelial, myoblast, and preadipocyte cell differentiated states. The results are summarized in Figure 6 for the genes *PCDHA4, PCDHB2*, and *PCDHGA12*. In the case of *PCDHA4* and *PCDHB2*, the differences in percent methylation between embryonic and adult cells was highly significant (p< 0.0001). In the case of *PCDHA4* and *PCDHB2*, DMRs tended to be associated with gene bodies rather than promoter regions (Supplementary Figure 3). The DMR associated with the adult-onset gene *PCDHGA12* overlapped with the promoter of the gene (Supplementary Figure 3), and was significantly (p< 0.001) hypermethylated in embryonic cells, despite the accessibility and expression of the gene being adult onset. We therefore conclude that the sarcoma cell lines studied exhibited a DMR pattern more closely resembling an embryonic pattern of methylation than that of their adult counterparts (Figure 6).

**Figure 6.**
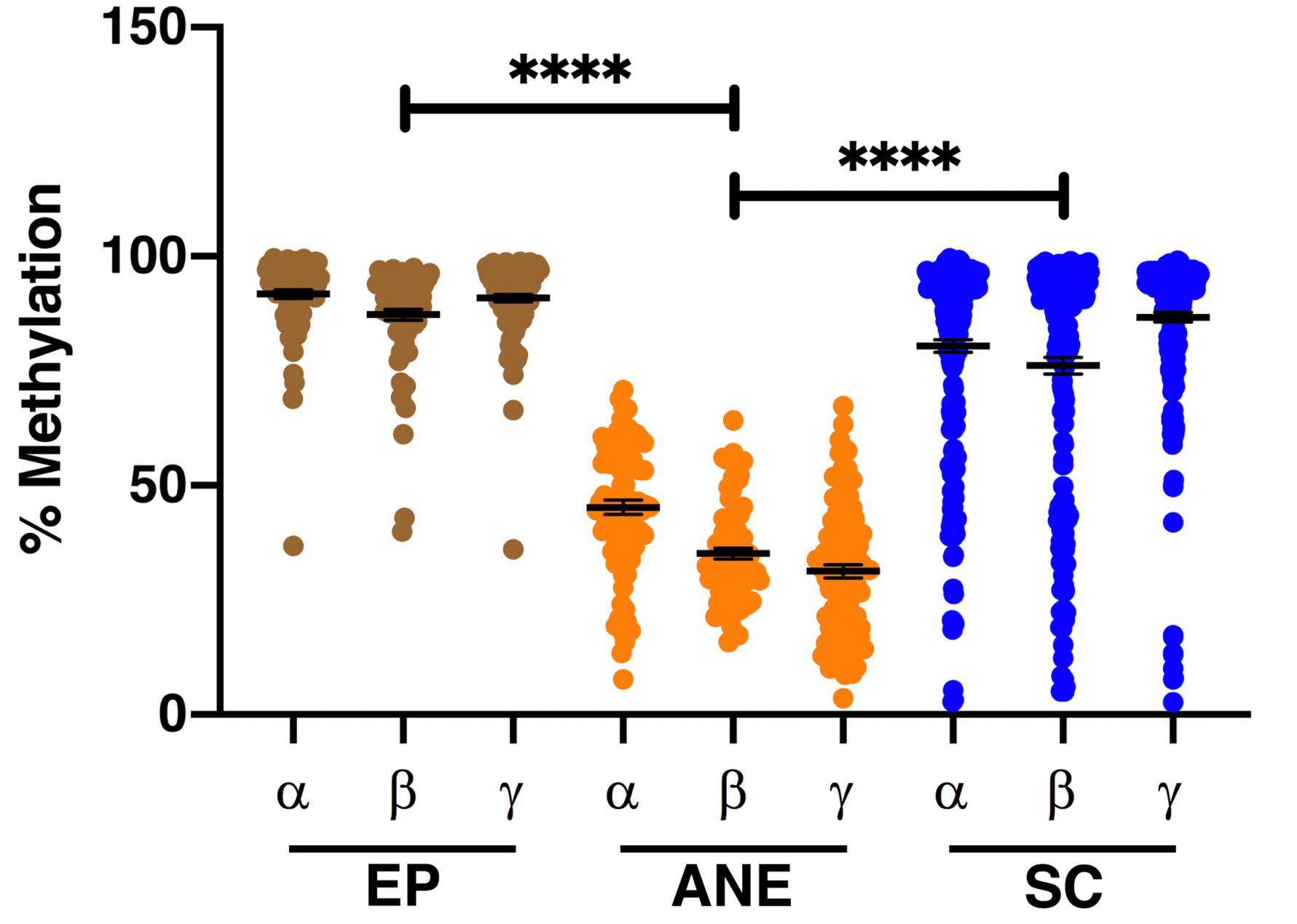
Percent methylation of representative significant DMRs within the CPL. Percent methylated CpGs is compiled for representative DMRs of the α, β, and γ clusters (*PCDHA4, PCDHB2,* and *PCDHGA12* respectively) for embryonic progenitor cells (EPs), Adult non-epithelial stromal and parenchymal cell types (ANE), and sarcoma cell lines (SC). Error bars S.E.M.

The CpG Island Methylator Phenotype (CIMP) is a commonly-studied category of DMRs hypermethylated in cancer^25^. A panel of CIMP DMRs markers commonly used in colorectal cancer include those associated with *CACNA1G*, *CDKN2A, CRABP1, IGF2*, *MLH1, NEUROG1*, *RUNX3*, and *SOCS1* genes^26^. As shown in Supplementary Figure 4A and B, while these DMRs within CpGIs show hypermethylation in a number of the cancer cell lines, they do not appear to be significantly (q-value < 0.05) differentially-methylated in embryonic vs adult cell types. The *IGF2* and *RUNX3* CpGIs were an exception, however they are significantly hypermethylated in adult cells, in contrast to the DMRs in the CPL locus. Therefore, we conclude that the DMRs associated with the embryo-onco phenotype in the CPL do not reflect the CIMP as it is commonly described, but instead may reflect an embryo-specific methylator phenotype which we will refer to as CIMP-E.

### Chromatin Architecture in the CPL Appears Altered in the Embryonic vs Adult Cell types in Alignment with Lamina-Associated Domains

As shown in Figure 5, the ∼0.6 MB region spanning the α and β clusters displayed markedly less accessibility in both adult osteogenic mesenchymal cells (MSCs) and adult aortic endothelial cell lines compared to their embryonic counterparts. Lamin interactions, such as those associated with lamin A/C and lamin B1 can impart heterochromatic alterations in chromatin spanning > 0.1 megabases associated with the nuclear periphery (lamin-associated domains (LADs)), we therefore examined the association of LADs with the CPL locus. Recognizing that lamin A appears to have dual roles at the nuclear periphery (isolated by sonication), and intranuclear (isolated by micrococcal nuclease digestion), we utilized previously identified lamina-interacting domains (LiDs) associated with lamin B1, and lamin A in HeLa cells from both sonicated and micrococcal nuclease-prepared samples assayed by ChIP-seq^27^ as a more comprehensive survey of chromatin interactions with lamins.

As shown in Figure 7, the lamin B1 LAD co-localized closely with the inaccessible region of the CPL locus in adult cells and was demarcated by CTCF binding sites^28^. Furthermore, the micrococcal nuclease-prepared Lamin A ChIP-seq sample appeared to closely overlap the region associated with Lamin B1 but to also extend over the γ locus and a 3’ superenhancer region identified using dbSuper^29^.

**Figure 7.**
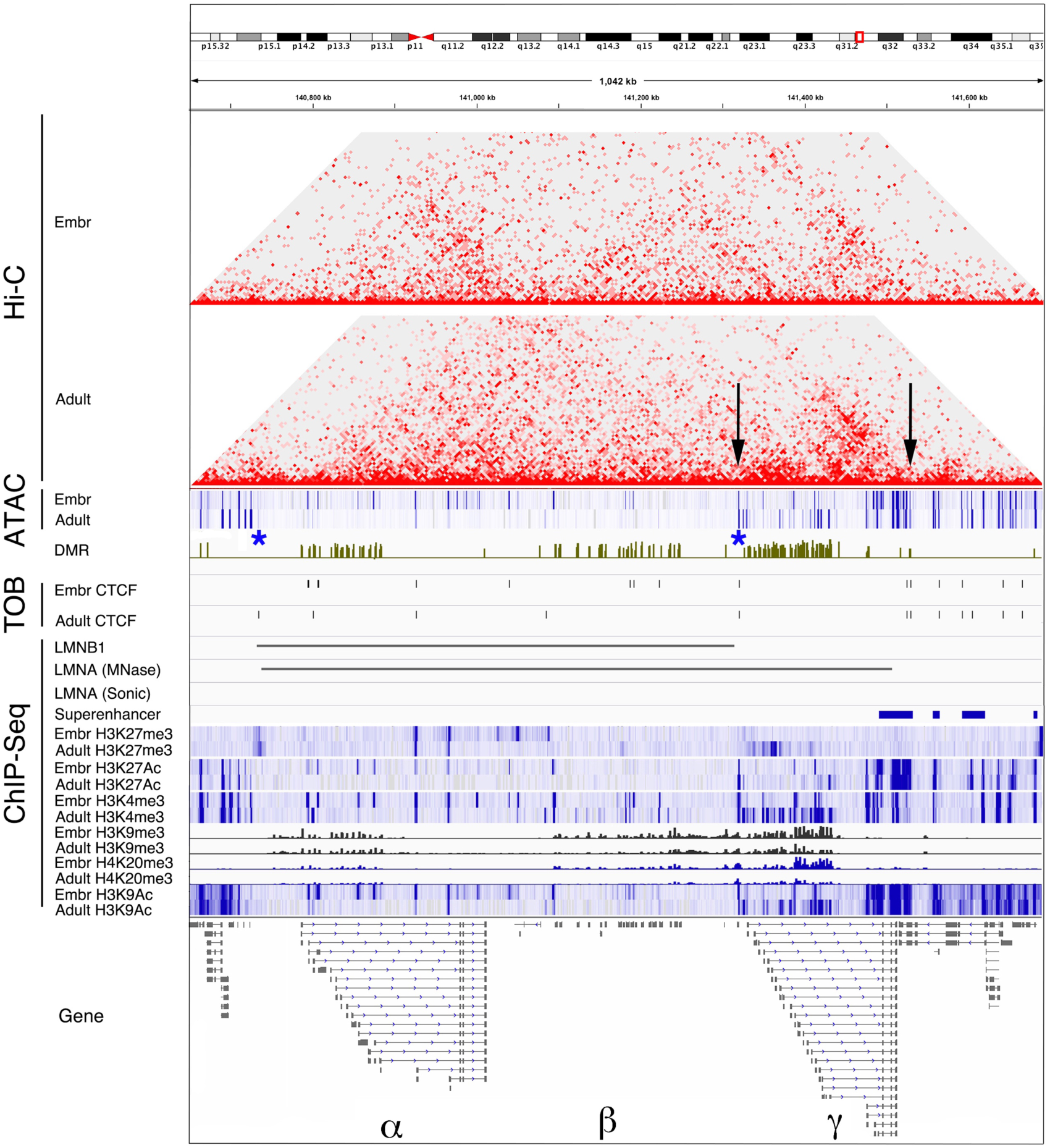
Chromatin architecture in the CPL. Rows represent: 1) chromatin cis-interactions determined by Hi-C analysis showing associations between the location of superenhancers in the CPL region with the promoters of the gamma locus of isoforms in adult cells (arrows); 2) ATAC-seq of the embryonic progenitor osteogenic mesenchymal line 4D20.8 (Embr Mesen) and adult-derived osteogenic mesenchyme (MSCs). Blue asterisks make region of apparent decreased accessibility in adult cells; 3) Differentially Methylated Regions (DMRs) are shown where elevated green represents hypermethylation in embryonic cells and depressed red signal represents relative hypomethylation in embryonic compared to adult cells; 4) CTCF binding sites in Embr Mesen and Adult Mesen based on TOBIAS analysis of ATAC footprints; 5) ChIP-seq values for CTCF; ChIP-seq results for LMNB1, LMNA (micrococcal nuclease treated (MNase), LMNA (sonicated); 6) location of superenhancers in the CPL region. 7) ChIP-seq data in paired embryonic and adult cells resulting from precipitation of chromatin with antibodies directed to H3K27Ac, H2K4me3, H2K9me3, and H4K20me3, and H3K9Ac for the entire CPL. (hg38 Chromosome position: Chr5: 140,650,000-141,700,000), image captured from IGV.

LADs have commonly been associated with peripheral heterochromatin marked by histone mark H3K9me3, demarcated by CTCF binding sites, which are thought to play a role in recruiting transcription factors as well as functioning as insulators in defined topological domains thereby regulating the function of enhancers, and for instituting a repressive environment for gene expression^28, 30^. Consistent with these markers of LADs, we observed a pronounced island of the heterochromatic marker H3K9me3 as well as H4K20me3 overlapping with the CPL LAD in both embryonic and adult cells. However, as described above, the CPL isoforms are nevertheless expressed in diverse embryonic and adult cell types despite these heterochromatic marks. In addition, although it is generally accepted that H3K4me3 marks are reduced in the repressive environment of a LAD^28^, we observed strong H3K4me3, as well as H3K27Ac, and H3K9Ac marks in association with all the expressed isoforms within the α, β, and γ loci (Figure 7 and Supplementary Figure 3). H3K27Ac and H3K9Ac, like H3K4me3 are considered markers of active gene expression^31^, consistent with the pattern of gene expression noted above.

ATAC-Seq showed open footprints for CTCF binding sites as determined by TOBIAS in regions other than those associated with CPL CpGIs (Figure 7, marked with black arrows). The most 3’ of these two CTCF sites is outside of the γ cluster and resides within a superenhancer region. The presence of CTCF binding at these two sites as determined by TOBIAS in all cells assayed (Figure 5), suggests that a potential topological domain exists in both embryonic and adult cells at this location. To confirm this potential constitutive topological domain, we utilized chromosome conformation capture (Hi-C) to reconstruct potential pairwise interactions based on domain topology. As shown in Figure 7, both embryonic and adult cells appear to show an interaction between the aforementioned twin CTCF sites (marked by the black arrows).

### Homologous Expression of CPL Isoforms in Varied Types of Differentiated Cells and Anatomical Locations

Homologous expression of CPL isoforms has been previously demonstrated to lead to cell-cell aggregation^13, 32^. While CPL isoforms are reported to be expressed stochastically from each allele in neuronal cells, presumably to regulate self-recognition^33^, patterns of CPL isoforms would be expected to be uniquely and uniformly expressed within particular differentiated cell types if they were to form homophilic interactions with their ectodomains and thereby play a role in embryonic tissue morphogenesis. We tested whether our data was consistent with this hypothesis *in silico* by examining the pattern of CPL isoform expression in biological replicates of particular differentiated cell types with hierarchical clustering objectively determining expression homology. We therefore performed a hierarchical clustering of a subset of the EP and ACs. As shown in Figure 8, ES cells (ESI-017 and ESI-053) clustered together but with the greatest divergence from the diverse differentiated cell types. The next layer of clustering effectively distinguished EP and ANE cells as predicted, since the basis of this study was their differential expression of α, β, and γ isoforms between these two categories of cells. Most significantly, replicates of embryonic ES-derived vascular endothelium (30-MV2-10 and 30-MV2-17) clustered closely together. Similarly, ES-derived progenitors of cartilage (4D20.8 and SK11)^34^, and resulting chondrocytes clustered closely together. In the case of adult cells, ectodermally-derived cell types such as keratinocytes, melanocytes, and astrocytes clustered closely as did dental pulp and skeletal myoblasts. Interestingly, Adult aortic endothelial and adult aortic smooth muscle cells reproducibly clustered together, consistent with their anatomical colocation and capacity for self-assembly^35^. Therefore, this *in silico* modelling is consistent with the hypothesis that combinations of CPL isoform expression in diverse somatic cell types may uniquely profile them, potentially playing a role in the adhesion of similar cells, or their adhesion to anatomically-related cells.

**Figure 8.**
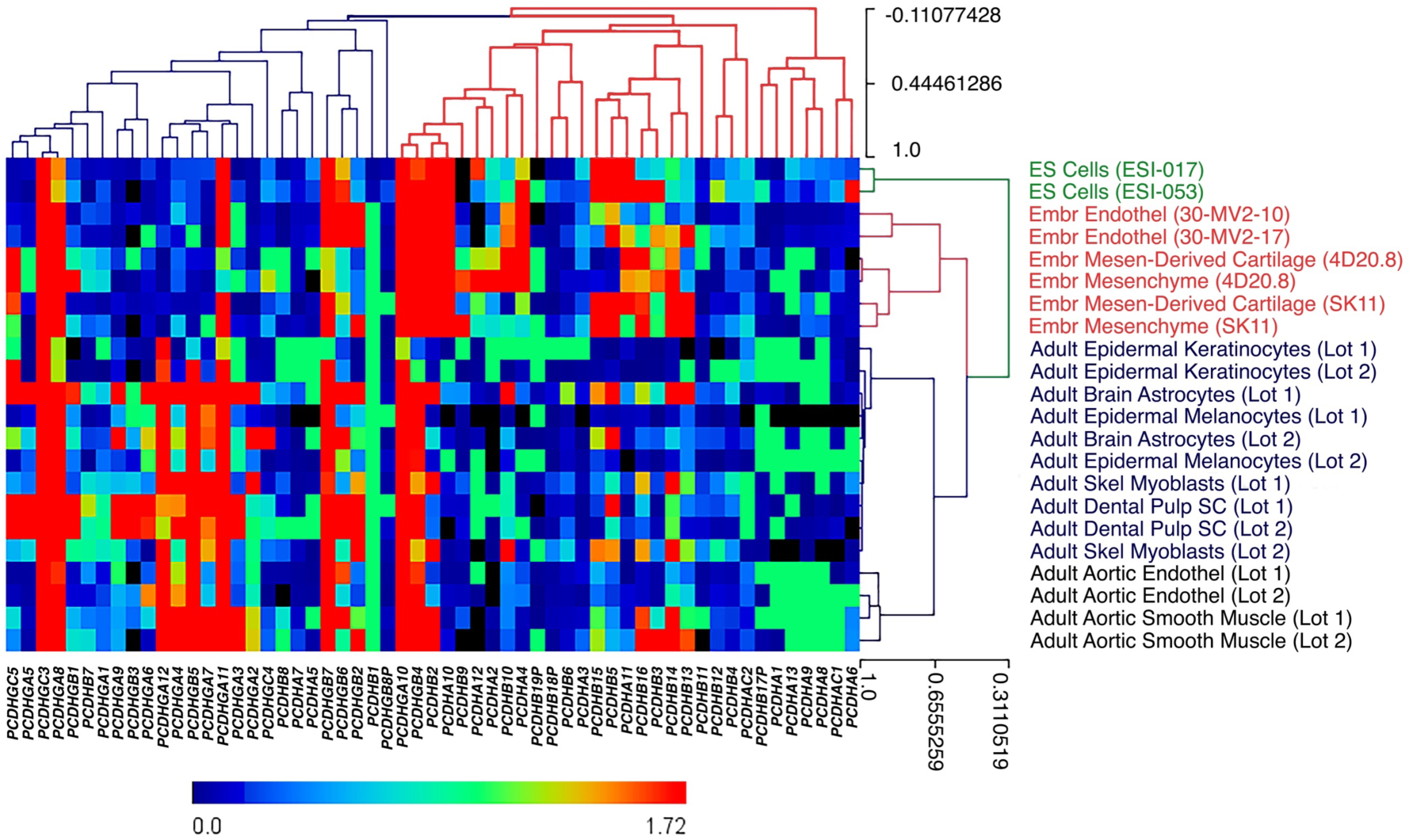
Heatmap of expression of all CPL genes in human ES cells, diverse embryonic progenitor cell types, and adult cells. RNA-seq FPKM values for isoform expression of every isoform of the α, β, and γ clusters in the CPL is heat mapped for select pluripotent stem cells (human ES cells), EP cell lines and adult cell types.

### CPL γ Isoforms are Down-Regulated During Cell Senescence *In Vitro*

The alterations we observed in α, β, and γ isoform gene expression in the course of embryonic-fetal development may reflect evolutionary selection for tumor suppression once embryonic organogenesis is complete. Since telomerase repression early in embryonic development leading to subsequent somatic cell replicative mortality is believed to reflect a similar example of antagonistic pleiotropy, we asked whether there are any alterations in CPL isoform expression occur during the aging of dermal fibroblasts *in vivo* and *in vitro*.

As shown in Supplementary Figure 5A, regardless of whether the γ isoform was down-regulated during the EFT as seen in *PCDHGB4*, or up-regulated (*PCDHGA12*), all γ isoforms showed down-regulation during senescence *in vitro* (Supplementary Figure 5B). In regard to *in vivo* aging, significant up-regulation of γ isoforms including *PCDHGB4* and *PCDHGA12* were observed in postnatal aged fibroblasts compared to fetal (synchronized dermal fibroblast lines from the medial aspect of the upper arm aged 11-83 years and 8-16 weeks gestation respectively) (Supplementary Figure 6A,B). However, no significant trends were observed during aging *in vivo* in the post-natal period. In addition, dermal fibroblasts derived from the Hutchinson-Gilford Progeria Syndrome (HGPS) showed no significant differences compared to 12 age-matched control lines (see Supplementary Figure 6 for the examples of *PCDHGB4* and *PCDHGA12*). However, as shown in Figure 9 and Supplementary Figure 6A,B, isoforms in the γ locus decreased significantly with replicative senescence *in vitro*. Culture of cells in log growth in 10% serum as opposed to quiescence in 0.5% serum appeared to increase *PCDHGA12* expression (Supplementary Figure 6B), highlighting the importance of growth and culture synchronization in expression analysis of CPL isoforms.

**Figure 9.**
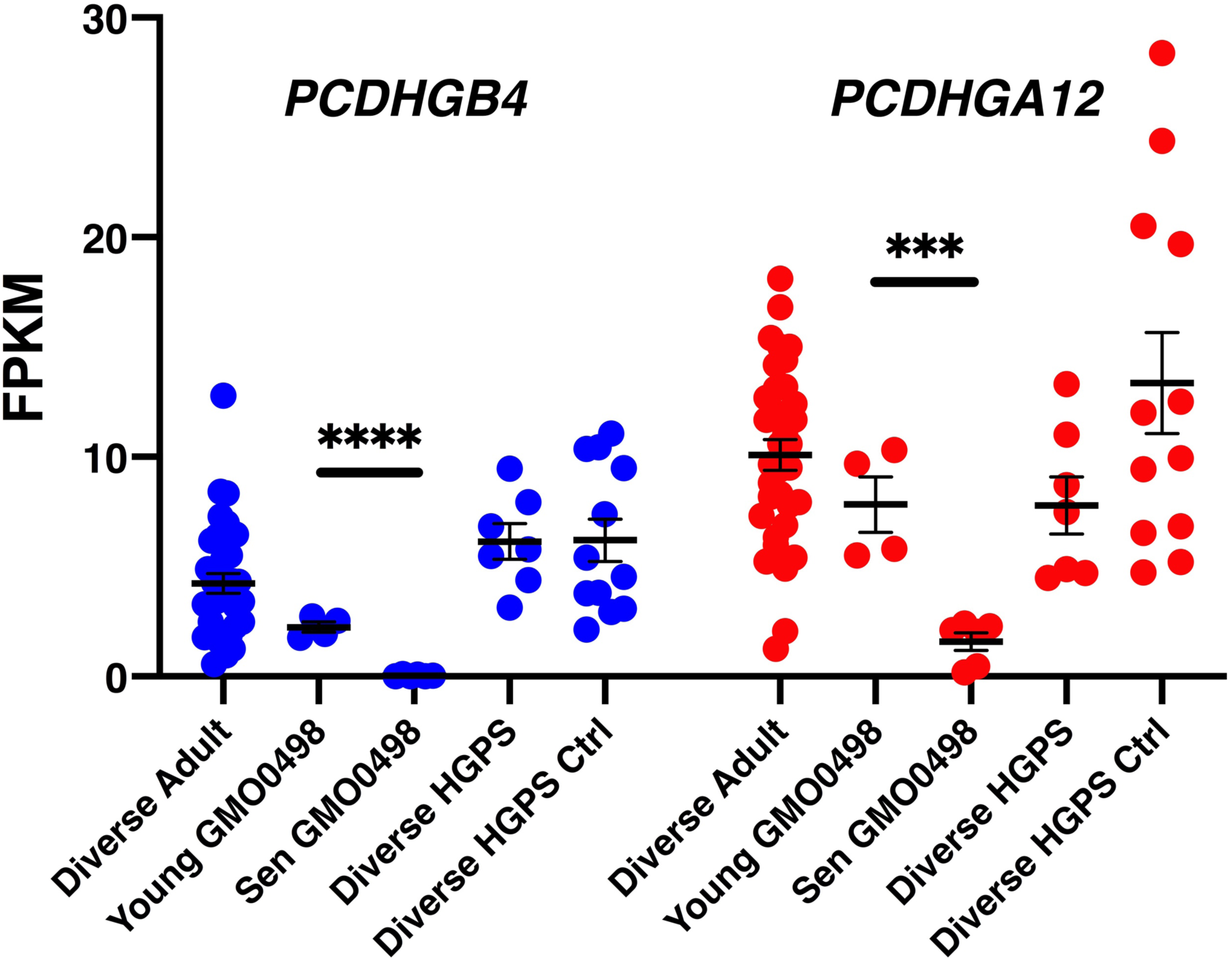
*PCDHGB4* and *PCDHGA12* gene expression during aging *in vitro* and in HGPS. RNA-seq FPKM values for *PCDHGB4* (blue) and *PCDHGA12* (red) isoform expression in synchronized diverse dermal adult fibroblasts (aged 11-83 years); young (P19) and senescent (P38) GM00498 (3 year-old donor) dermal fibroblasts, dermal fibroblasts from Hutchinson-Gilford progeria syndrome (HGPS) patients and age-matched controls. (*** p<0.001) (**** p< 0.0001) Error bars represent mean and standard error of the mean.

### Correlation of lamin and CPL isoform expression in development, aging, and cancer

We next asked whether CPL isoform expression correlated with that of lamin A or lamin B1 during development, aging, or cancer. As previously reported^36^, we observed that pluripotent stem cells expressed markedly lower levels of *LMNA*, while relatively high levels of *LMNB1* compared to fetal or adult counterparts (Supplementary Figure 7A,B). In regard to *in vitro* senescence, we again observed serum and/or growth-related effects on expression of both lamins. When comparing early passage (P19) dermal fibroblasts with senescent counterparts (P38), we saw a significant increase in *LMNA* and a significant decrease in *LMNB1* as previously reported^37^.

In the case of the diverse sarcoma and carcinoma cell lines used in this study, we saw a significant trend toward increasing expression of *LMNA* and decreasing expression of *LMNB1* correlating with *PCDHGA12* expression in the cancer cell lines (Supplementary Figures 8A,B). The role of lamins A and B1 in organizing expression domains leads to the intriguing question as to whether they play causative role in the regulation of altered gene expression in the embryonic-fetal transition during normal development and cancer or are downstream of other yet unknown regulatory events.

## Discussion

In numerous vertebrate species, the transition from the embryonic/larval stages of development to that of the fetus or adult is characterized by a loss of cell-cell recognition and epimorphic tissue regeneration in many somatic tissues^38^. Rigorous interrogation of the molecular pathways regulating this transition in humans is currently limited by the lack of a robust and reproducible *in vitro* model for the first eight weeks of development. We therefore employed a novel *in vitro* system wherein diverse hES cell-derived clonal embryonic progenitors were compared to fetal and adult-derived counterparts. Clonal EP cells show significant stability with passaging, thereby enabling reproducible experimentation^23, 39^. We therefore conclude that this model may provide a means to test hypotheses regarding the EFT and thereby glean insights into the molecular biology of regeneration.

William Dreyer proposed that a complex repertoire of cell adhesion molecules regulated selective cell-cell adhesion during animal development^10^. The clustered protocadherin locus provided one such source of diversity, however, research to date has largely focused on the role of stochastic expression of the genes in neuronal self/non-self recognition in trans, thought to play an important role in neural network formation^15, 40^. Consistent with this role in neural development, we found that clonal isolates of iPSC-derived neural cells *in vitro* displayed diverse patterns of α/γ isoform expression. Moreover, consistent with our results, these cultured pluripotent stem cell-derived neural cells failed to transition an adult-like CPL phenotype when maintained *in vitro*, or even when transplanted *in vivo*^40^.

As described in Aron Moscona’s studies of the reassociation of chick embryonic anlagen such as chondrogenic mesenchyme and skeletal muscle cells^6^, pre-EFT tissues display a remarkable capacity for scarless regeneration following trauma. We therefore suggest that the CPL locus provides a complex organizing center for diverse differentiated somatic cell type-specific adhesion molecules as theorized by Dreyer.

In this study, we extend the analysis of CPL isoform expression to diverse somatic cell types outside of the CNS. We detected discrete isoforms from the α and β clustered protocadherin locus in EP cells (ES-derived clonal embryonic progenitor lines with negligible fetal or adult-specific *COX7A1* expression)^19^. The α and β CPL expression was comparable or greater than cultured neuronal cell types. In contrast, most adult-derived stromal, parenchymal, and epithelial cell types studied showed reduced or non-existent expression of isoforms in the α and β clusters. In the case of the γ locus, some genes such as *PCDHGB4* and *PCDHGB6* showed embryonic up-regulation, while others like *PCDHGB5* and *PCDHGA12* showed negligible expression in pluripotent stem cells and embryonic progenitors, but up-regulation in most adult differentiated cell types. It is surprising that these expression patterns in the CPL in non-neuronal cell types have not previously been reported. One explanation is that transcriptomic alterations in the α and γ clusters require isoform registration not commonly patterned in microarrays, and therefore they were largely undetectable before the widespread use of RNA-sequencing.

The timing of the EFT in early passage cultures of dermal fibroblasts was previously assayed using *COX7A1* expression, a robust marker of EFT in stromal cell types^19^. We reported that *COX7A1* expression was essentially undetectable in pluripotent stem cells and their embryonic progenitor derivatives, but low levels of expression began to be detectable in early passage cultures derived from fetal skin as early as eight weeks of gestation. In this study, we observed that the fetal-adult pattern of CPL gene expression was largely complete by eight weeks (with the possible exception of the embryonic-specific gene *PCDHA4* where FCs showed a small but significant elevation over adult counterparts). Therefore it would seem reasonable to conclude that the transition from embryonic to fetal CPL isoform expression occurs before eight weeks of gestation. Further studies in human and rodent models may shed light on the temporal as well as anatomical staging of CPL isoform transitions.

Altered expression of CPL genes in cancer has occasionally been reported previously. For example, a bioinformatics-based search for pan-cancer markers found both *COX7A1* and *PCDHA12* (both being adult-onset genes), as being down-regulated in 10 carcinoma and adenocarcinoma types compared to normal tissues^41^. We extended that observation to inlcude sarcomas to enable a comparison with their corresponding differentiated cell types, i.e., osteosarcomas compared to embryonic and adult osteogenic mesenchyme, and comparable pairing with myoblasts and white preadipocytes. We observed a striking shift to and embryonic pattern of CPL isoform expression in the majority of cancer lines. For example, in the case of *PCDHGA12*, 21/39 (54%) of sarcoma lines and 33/35 (94%) of carcinoma and adenocarcinoma cell lines did not express the gene at a level of at least 0.5 FPKM, whereas 90/92 (98%) of adult non-epithelial cell types did. The correlation of the embryonic phenotype in various cancer cell lines (sarcoma and carcinoma lines from varied tissues of origin) with an absence of *COX7A1* expression, suggests that a large percentage of cancer cell lines may display an embryonic phenotype in regard to the expression of genes beyond those of the CPL.

As early as 1859, based on histological comparisons, Rudolf Virchow proposed that cancers originated “by the same law that regulated the embryonic development.^42^” Cancer cells and early embryonic cell types have long been recognized to share certain properties including an increased dependance on aerobic glycolysis (the Warburg Effect)^43^. Indeed, it has been proposed that cancer, or cancer stem cells in general, are a result of failed embryonic cell maturation or reprogramming of mature cells to an embryonic state^44^. However, since we observed a heterogeneity in the shift to embryonic CPL expression, further study will be necessary to determine the significance of embryonic gene expression in some cancer cells and adult-like expression in others. Such a comparison may expand our understanding of what is currently regarded as “cancer stem cells”, with potential implications for cancer prognosis and therapy.

It is intriguing that the majority of sarcoma, carcinoma, and adenocarcinoma cell lines tested simultaneously exhibited an embryonic pattern of CPL gene expression as well as embryonic DMR methylation. Altered DNA methylation in aging and cancer is increasingly a subject of intense interest. In the case of cancer, hypermethylated CpGIs are frequently associated with the repression of tumor suppressor genes, an example being the CpG Island Methylator Phenotype (CIMP) originally described in colon cancer^25^. One such CIMP marker is the tumor suppressor locus of P16. The PCDH β isoforms have been reported to be hypermethylated in neuroblastoma and reported to allow a modified CIMP that predicts poor outcome^45, 46^. More recently, hypermethylation of CPL genes has been reported in numerous tumor types^47^. Using the classical CIMP DMRs: *CACNA1G*, *CDKN2A, CRABP1, IGF2*, *MLH1, NEUROG1*, *RUNX3*, and *SOCS1,* we saw no correlation with the embryo-onco phenotype, indeed, in the case of the genes *IGF2* and *RUNX3*, CPGIs were detected that were hypermethylated in adult as opposed to embryonic cells. It therefore seems reasonable to conclude that the hypermethylation of the CPL isoforms in cancer is not associated with CIMP, but instead may reflect an independent set of methylation marks we designate CIMP-E. The pan-tissue nature of the EFT may lead to a pan-cancer CIMP-E profile consistent with a global developmental shift in somatic cells reminiscent of global *TERT* repression in numerous somatic cells once the associated developmental stage is completed^48^.

The CPL exhibits a number of unique features that lead us to believe it would provide a useful model system for the study of developmental alterations in gene expression and chromatin structure dynamics. First, the locus overlaps with Lamin A and B1 LADs and was demarcated by CTCF binding sites. Both lamins are differentially-expressed during development. *LMNB1* is expressed at the highest levels in pluripotent stem cells, shows decreased expression in fetal skin fibroblasts as a function of age of gestation, and decreases further during *in vitro* senescence. *LMNA* is expressed a low levels in pluripotent cells, shows increased expression in most somatic cell types (an exception being neurons), and on a protein level, levels increase during embryogenesis and fetal development at varied timepoints depending on cell type (∼E14-15 in mouse dermal fibroblasts)^49, 50^. It is therefore important to determine the role that these lamins, in particular, the regulation of protein levels of prelamin A vs mature lamin A have in regulating not only chromatin structure, but also EFT-related alterations in gene expression.

A second interesting feature of the CPL locus is the pronounced H3K9me3 histone modification spanning the locus may fall within the definition of a large organized chromatin lysine (K) domain (LOCK). H3K9me3-rich LOCKs are generally associated with centromeric and telomeric DNA^51^, or LADs^52^. On chromosome 5, ChIP-seq revealed three pronounced H3K9me3 LOCKS: subtelomeric and pericentromeric DNA, and the CPL locus (data not shown). LOCKs comprised of H3K9me3-rich regions are proposed to stabilize the differentiated state of somatic cells, hence the designation “differentiation bound regions” (DBRs). Further research may help elucidate the role of these LOCKs in development and/or in preventing illegitimate recombination in repeat regions, such as the isoform repeats of the CPL.

A third attractive feature of the CPL is the observed down-regulation of γ isoform expression with *in vitro* senescence. *LMNB1* expression has been shown to be markedly down-regulated with *in vitro* aging. In addition, methylated DNA immunoprecipitation from young and aged monocytes showed the γ locus to be hypermethylated with age^53^. It is well-established that both *in vitro* and *in vivo* aging of cells is associated with altered CpG methylation that can be assembled into reliable age classifier^54^. Taken together, these observations suggest that the CPL may provide a single model to further define the relationship between developmental transitions, such as those associated with the EFT and aging, nuclear architecture, and epigenetic markers such as DNA methylation markers of aging (AGEm).

It is well-documented that spatiotemporal alterations in chromatin architecture occurs during pluripotent stem cell differentiation^55^ however, potential changes associated with the EFT have received little if any attention to date. The striking co-localization of ATAC inaccessibility and ChIP-seq localization of lamin B together with the lamin A-specific association with the gamma locus hints at a potential regulatory role of these lamins in CPL isoform expression. Figure 10 illustrates a proposed model of the varied chromatin architecture and resultant gene expression in the locus in response altered lamin interactions during the transition to the non-regenerative fetal-adult state and the re-emergence of the embryonic phenotype in cancer. The progressive loss of lamin B1 and increasing levels of lamin A mRNA (and potentially more importantly, mature lamin A protein), leads to a reorganization of chromatin structure with the loss of an embryonic CPL code in the particular differentiated cell type. We propose that the loss of this embryonic CPL code plays a role in the loss of regenerative potential. The increased expression of γ isoforms post-EFT may also play a role in tumor suppression. The γ isoforms have been implicated, for instance, in regulating canonical wnt signaling^56^. More research will be necessary to uncover the potential effects of these alterations in the growth and metastasis of cancer cells.

**Figure 10.**
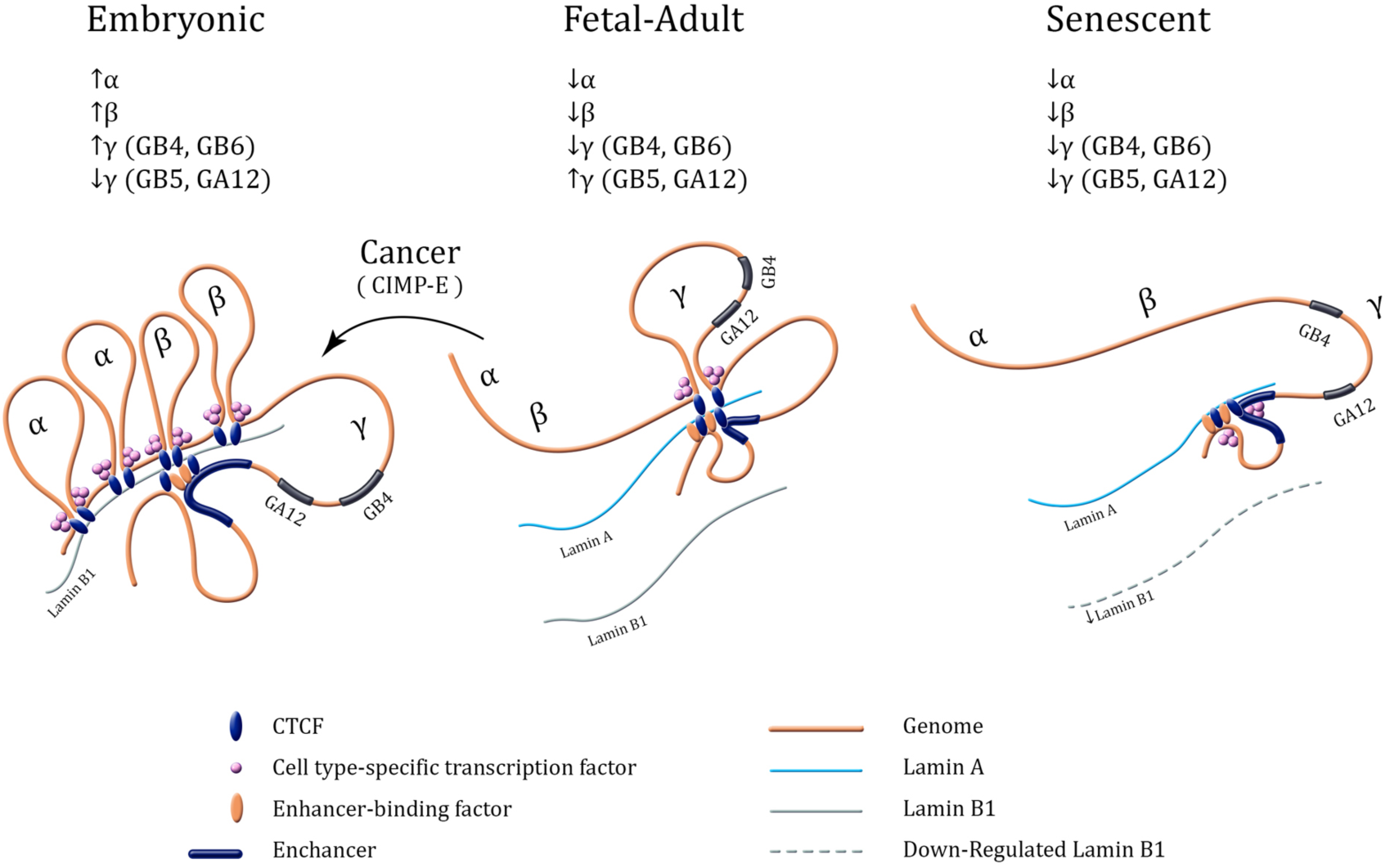
Model of potential transition in chromatin architecture during development, cellular aging, and cancer. During embryogenesis (up to approximately Carnegie Stage 23 of mammalian development), Lamin B1 predominates and facilitates CTCF binding, topological domains, and expression of cell type-specific CPL isoforms from the α and β clusters. Once organogenesis is complete, up-regulation of Lamin A facilitates an alteration in topological domains such that only isoforms from the γ cluster are expressed. In replicative senescence, *LMNB1* expression is down-regulated together with expression of genes in the γ cluster. The majority of cancer cell lines show a CpG Island Methylator Phenotype (CIMP) in the CPL locus designated herein as CIMP-E.

Assuming the CPL plays a role in embryonic development, and therefore, in epimorphic regeneration, strategies to reverse developmental aging and induce a regenerative state in context of disease, may focus on means of restoring the locus to an embryonic pattern of gene expression. These strategies called induced tissue regeneration (iTR), or partial reprogramming may, for instance, employ the transient administration of transcription factors that have the potential to revert fully-differentiated cells to a pluripotent state^38, 57^. Reprogramming by somatic cell nuclear transfer or by the exogenous administration of transcriptional regulators is typically an inefficient process^58, 59^, suggesting barriers to the access of chromatin. Chromatin modified by H3K9me3 histone modification is enriched at constitutively heterochromatic regions such as telomeric and centromeric chromatin. Therefore, these regions would be expected to be relatively inaccessible to pioneer reprogramming factors like KLF4, POU5F2 (OCT4), and SOX2. Outside of the zones of constitutive heterochromatin, H3K9me3 modification may play a role in stabilizing differentiated states in somatic cells, regions designated differentiated bound regions (DBRs)^60^. Reduction of H3K9me3 through the targeted down-regulation of the methyltransferases *SUV39H1, SUV39H2*, and *SETDB1*^60^ has been shown to increase the efficiency of induced pluripotency. Therefore, in the context of the transient expression of reprogramming factors to induce tissue regeneration, the selected removal of H3K9me3 marks at the CPL may increase the frequency of reverting aged somatic cells to an embryonic pattern of CPL expression, thereby improving the regenerative response.

The further elucidation of the potential role of CPL isoforms in embryonic development, like the role uncovered for the genes in neuronal function, may provide important new insights into pluripotent stem cell-derived organoid formation, the loss of regeneration and initiation of scarring in fetal and adult tissues, induced tissue regeneration, and perhaps even new insights into aspects of oncogenesis such as the propensity for discrete tumor formation as opposed to dispersed metastasis.

## Supporting information

Differential Expression of Clustered PCDH-Supplementary Materials

## Acknowledgements

We wish to thank Irina Turovets of Biotech Graphic Design for artistic contribution in Figure 10. We also acknowledge high-quality sequencing service from the Beijing Genomics Institute (BGI) for their assistance in performing RNA-seq and WGBS of our samples. Lastly, we’d like to thank Advanced Bioscience Resources for their help in providing tissues used in the study.

## Competing Interests Statement

HS, IL, JL, PS, JJ, DL and MDW are employees of and stockholders in AgeX, Inc., (Alameda, CA). KBC is an employee of Eclipse Innovations, (San Diego, CA) and a stockholder of AgeX, Inc.

## Methods

### Cell Culture

Pluripotent stem cells (PSCs) (hESCs and iPSCs) were cultured on Matrigel in mTeSR1 medium in a humidified incubator at 37°C with 5% O_2_ and 10% CO_2_, Embryonic Progenitor Cells (PCs) were cultured on 0.1% gelatin in their specific growth medium used originally when cloning and scaling the line in a humidified incubator at 37°C with 5% O_2_ and 10% CO_2_. The EP cell lines 4D20.8 and were cultured in DMEM 20% FBS and PromoCell endothelial (MV2) growth medium respectively. Cells were routinely passaged 1:3 at or near confluence using 0.05% trypsin. Adult-derived cells were cultured in their respective optimal growth mediums on 0.1% gelatin in Corning cultureware at 37°C with 5% O_2_ and 10% CO_2_ or in select cases, total RNA was purchased from ScienCell (Carlsbad CA). The cancer cell lines were obtained from ATCC and cultured as suggested by ATCC prior to harvest. Prior to analysis, all lines were synchronized in quiescence when possible by culturing five days in 10% of the normal serum or related growth factor supplements.

### RNA isolation

RNA was prepared upon lysis with RLT with 1% 2-βME, using Qiagen RNeasy mini kits (Cat#74104) following manufacturer’s directions. The extracted RNA was then quantitated using a NanoDrop (ND-1000) spectrophotometer and the labeled tubes were stored at -80°C for later use.

### RNA-sequencing

Library Construction was performed by using Illumina Truseq mRNA library prep kit following manufacturer’s directions. Library QC and library pooling was accomplished using Agilent Technologies 2100 Bioanalyzer to assay the library fragments. qPCR was used to quantify the libraries. Libraries were pooled, which have different barcodes/indexing and sequencing, in one lane. The paired-end sequencing was performed using the Illumina HiSeq4000 sequencing instrument, yielding 100-bp paired-end reads. The sequencing was performed by BGI AMERICAS CORPORATION.

### RNA-Seq Data Analysis

The fastq files containing a minimum of 25 million reads per sample obtained by sequencing were analyzed using the Tuxedo protocol^61^. Reads are aligned against GRCh38 using short read aligner Bowtie2 (release 2.2.7) within the TopHat (release 2.1.1) splice junction mapper. Bowtie2 indices, as well as GRCh38 genome annotation) were obtained from Illumina, Inc. iGenomes. Alignment files were assembled into transcripts, and the abundances estimated using cufflinks 2.2.1 release. To allow for high abundance of transcripts, the parameter –max bundle-frags 2000000 was used. Cufflinks gtf files were merged with genes.gtf of GRCh38 annotation using Cuffmerge (release 2.2.1) into unified transcript catalog. Transcript abundance levels were computed using Cuffquant, and the resulting data normalized using Cuffnorm (both 2.2.1 Cufflinks release).

### Volcano Plot

Data analysis of the transcription levels (FPKM values) was carried out in R. Data was filtered to remove genes from the Y chromosome. FPKM values were rounded to two decimal places and low-expressing entities were removed by filtering for a mean FPKM value > 0.5 in either the embryonic or adult group and a mean > 0 in both groups. P-values were generated using a t-test with Bonferroni correction. Plots were generated in R.

### Statistical Analysis

Statistical significance of differences in FPKM values was determined using a two-tailed T-test assuming two samples of equal variance (homoscedastic). Error bars designate standard error of the mean unless otherwise noted.

### ATAC-seq

Embryonic progenitor cell lines (4D20.8 and 30-MV2-6), adult cell lines (MSC and HAEC), PSC lines, and cancer lines were harvested and frozen in culture media containing FBS and 5% DMSO. Cryopreserved cells were sent to Active Motif to perform the ATAC-seq assay. The cells were then thawed in a 37°C water bath, pelleted, washed with cold PBS, and tagmented as previously described^21, 62^. Briefly, cell pellets were resuspended in lysis buffer, pelleted, and tagmented using the enzyme and buffer provided in the Nextera Library Prep Kit (Illumina). Tagmented DNA was then purified using the MinElute PCR purification kit (Qiagen), amplified with 10 cycles of PCR, and purified using Agencourt AMPure SPRI beads (Beckman Coulter). Resulting material was quantified using the KAPA Library Quantification Kit for Illumina platforms (KAPA Biosystems), and sequenced with PE42 sequencing on the NextSeq 500 sequencer (Illumina).

Analysis of ATAC-seq data was very similar to the analysis of ChIP-Seq data (see below). Reads were aligned using the BWA algorithm (mem mode; default settings). Duplicate reads were removed, only reads mapping as matched pairs and only uniquely mapped reads (mapping quality >= 1) were used for further analysis. Alignments were extended in silico at their 3’-ends to a length of 200 bp and assigned to 32-nt bins along the genome. The resulting histograms (genomic “signal maps”) were stored in bigWig files. Peaks were identified using the MACS 2.1.0 algorithm at a cutoff of p-value 1e-7, without control file, and with the –nomodel option. Peaks that were on the ENCODE blacklist of known false ChIP-Seq peaks were removed. Signal maps and peak locations were used as input data to Active Motifs proprietary analysis program, which creates Excel tables containing detailed information on sample comparison, peak metrics, peak locations and gene annotations.

### TOBIAS analysis

To qualify the potential bindings of the transcription factors to the open regions characterized by ATAC-Seq, we used the TOBIAS algorithm^63^. Histogram peaks were characterized from the BAM files using MACS2 callpeak function of MACS^64^. To compensate tor Tn5 transposase site insertion bias, TOBIAS ATACorrect function was used, and the data stored in bigwig output files. Corrected bigwig files were characterized for peaks using MACS2 call peaks function with the following parameters: --shift -100 –extsize 200 –broad. Obtained peaks were searched for the transcription factor binding footprints using TOBIAS Footprint Scores function. Binding site footprint scores were characterized for transcription factor binding events with TOBIAS BINDetect for the pairs of samples (4D20 vs HMSC, and 30MV2-6 vs HAEC) against JASPAR 2020 database^65^. Resulting bed files were visualized by IGV^66^.

### Chromatin Immunoprecipitation

Cells were fixed with 1% formaldehyde for 15 min and quenched with 0.125 M glycine. Frozen cell pellets were sent to Active Motif to perform the ChIP-Seq assay. Chromatin was isolated by the addition of lysis buffer, followed by disruption with a Dounce homogenizer. Lysates were sonicated and the DNA sheared to an average length of 300-500 bp. Genomic DNA (Input) was prepared by treating aliquots of chromatin with RNase, proteinase K and heat for de-crosslinking, followed by ethanol precipitation. Pellets were resuspended and the resulting DNA was quantified on a NanoDrop spectrophotometer. Extrapolation to the original chromatin volume allowed quantitation of the total chromatin yield.

An aliquot of chromatin (30 μg) was precleared with protein A or G agarose beads (Invitrogen). Genomic DNA regions of interest were isolated using 4 ug of antibody against the specific histone modification of interest. Complexes were washed, eluted from the beads with SDS buffer, and subjected to RNase and proteinase K treatment. Crosslinks were reversed by incubation overnight at 65° C, and ChIP DNA was purified by phenol-chloroform extraction and ethanol precipitation.

Quantitative PCR (QPCR) reactions were carried out in triplicate on specific genomic regions using SYBR Green Supermix (Bio-Rad). The resulting signals were normalized for primer efficiency by carrying out QPCR for each primer pair using Input DNA.

### ChIP Sequencing

Illumina sequencing libraries were prepared from the ChIP and Input DNAs by the standard consecutive enzymatic steps of end-polishing, dA-addition, and adaptor ligation. Steps were performed on an automated system (Apollo 342, Wafergen Biosystems/Takara). After a final PCR amplification step, the resulting DNA libraries were quantified and sequenced on Illumina’s NextSeq 500 (75 nt reads, single end). Reads were aligned to the human genome (hg38) using the BWA algorithm (default settings). Duplicate reads were removed and only uniquely mapped reads (mapping quality >= 25) were used for further analysis. Alignments were extended in silico at their 3’-ends to a length of 200 bp, which is the average genomic fragment length in the size-selected library, and assigned to 32-nt bins along the genome. The resulting histograms (genomic “signal maps”) were stored in bigWig files. For active histone marks, peak locations were determined using the MACS^62^ algorithm (v2.1.0) with a cutoff of p-value = 1e-7. For repressive histone marks or histone modifications with broad distribution, enriched regions were identified using the SICER^74^ algorithm at a cutoff of FDR 1E-10 and a max gap parameter of 600 bp. Peaks that were on the ENCODE blacklist of known false ChIP-Seq peaks were removed. Signal maps and peak locations were used as input data to Active Motifs proprietary analysis program, which creates Excel tables containing detailed information on sample comparison, peak metrics, peak locations and gene annotations.

### Whole-Genome Bisulfite Sequencing

DNA was prepared from cell lines using Qiagen DNeasy Blood and Tissue kits (Cat#69504) following manufacturer’s instructions. The labeled microfuge tubes were then stored at -80°C. Samples were sequenced as 150-bp paired-end reads, to the level of about 90G of raw data per sample. Sequencing was performed by BGI AMERICAS CORPORATION.

### WGBS data analysis

Whole genome bisulfite sequencing data analysis was performed on obtained fastq reads files using Bismark suite with default parameters^67^.

### Identification of DMRs

DMRs were identified from the WGBS data using Metilene software^68^. DMRs were defined with a q-value < 0.01 and a mean methylation difference > 0.15 in a window of at least 250 nt, eight CpGs, and signal in at least three of four of the embryonic or adult cell lines in the cohort.

### TAD Method

Chromatin conformation capture data was generated using a Phase Genomics (Seattle, WA) Proximo Hi-C 2.0 Kit, which is a commercially available version of the Hi-C protocol^69^. Following the manufacturer’s instructions for the kit, intact cells from two samples were crosslinked using a formaldehyde solution, digested using the Sau3AI restriction enzyme, and proximity ligated with biotinylated nucleotides to create chimeric molecules composed of fragments from different regions of the genome that were physically proximal in vivo, but not necessarily genomically proximal. Continuing with the manufacturer’s protocol, molecules were pulled down with streptavidin beads and processed into an Illumina-compatible sequencing library. Sequencing was performed on an Illumina HiSeq 4000.

Reads were aligned to the Homo sapiens genome assembly GRCh38 (hg38), also following the manufacturer’s recommendations. Briefly, reads were aligned using BWA-MEM^70^ with the -5SP and -t 8 options specified, and all other options default. SAMBLASTER^71^ was used to flag PCR duplicates, which were later excluded from analysis. Alignments were then filtered with samtools^72^ using the -F 2304 filtering flag to remove non-primary and secondary alignments. TopDom^73^ was used to identify topologically associated domains (TADs) at a 50kb resolution by computing the average contact frequency among pairs of chromatin regions (one upstream and one downstream) in a small window (w=5) around the 50kb bin. The resulting curve was used to identify local minima, or regions of low chromatin contact, along each chromosome. Regions of interest were plotted using pyplot and patches from the matplotlib Python package^74^. The BED files documenting TAD calls by TopDom were simplified and viewed as tracks in the USCS genome browser. Data visualization of obtained .hic files were visualized using the Juicebox suite^75^.

## References

1. Grunwald, G.B. The conceptual and experimental foundations of vertebrate embryonic cell adhesion research. Dev Biol (N Y 1985) 7, 129–158 (1991).

2. Steinberg, M.S. & Gilbert, S.F. Townes and Holtfreter (1955): directed movements and selective adhesion of embryonic amphibian cells. J Exp Zool A Comp Exp Biol 301, 701–706 (2004).

3. Wilson, H.V. On Some Phenomena of Coalescence and Regeneration in Sponges. J. Exp. Zool. 5, 245–258 (1907).

4. Spemann, H. & Mangold, H. Induction of embryonic primordia by implantation of organizers from a different species. 1923. Int J Dev Biol 45, 13–38 (2001).

5. Moscona, A. & Moscona, H. The dissociation and aggregation of cells from organ rudiments of the early chick embryo. J Anat 86, 287–301 (1952).

6. Li, G.X., Moscona, M.H. & Moscona, A.A. Effect of embryonic age on aggregability, histogenesis and biochemical differentiation in the embryonic chick and quail neural retina. Sci Sin B 27, 371–379 (1984).

7. Williams, G. Pleiotropy, Natural Selection, and the Evolution of Senescence. Evolution 11, 398–411 (1957).

8. Kim, N.W. et al. Specific association of human telomerase activity with immortal cells and cancer. Science 266, 2011–2015 (1994).

9. Dreyer, W.J.G., W.R.; Hood, L. The Genetic, Molecular, and Cellular Basis of Antibody Formation: Some Facts and a Unifying Hypothesis. Cold Spring Harb Symp Quant Biol 32, 353–367 (1967).

10. Dreyer, W.J. & Roman-Dreyer, J. Cell-surface area codes: mobile-element related gene switches generate precise and heritable cell-surface displays of address molecules that are used for constructing embryos. Genetica 107, 249–259 (1999).

11. Wu, Q. & Maniatis, T. A striking organization of a large family of human neural cadherin-like cell adhesion genes. Cell 97, 779–790 (1999).

12. Thu, C.A. et al. Single-cell identity generated by combinatorial homophilic interactions between alpha, beta, and gamma protocadherins. Cell 158, 1045–1059 (2014).

13. Schreiner, D. & Weiner, J.A. Combinatorial homophilic interaction between gamma-protocadherin multimers greatly expands the molecular diversity of cell adhesion. Proc Natl Acad Sci U S A 107, 14893–14898 (2010).

14. Kaneko, R. et al. Allelic gene regulation of Pcdh-alpha and Pcdh-gamma clusters involving both monoallelic and biallelic expression in single Purkinje cells. J Biol Chem 281, 30551–30560 (2006).

15. Lefebvre, J.L., Kostadinov, D., Chen, W.V., Maniatis, T. & Sanes, J.R. Protocadherins mediate dendritic self-avoidance in the mammalian nervous system. Nature 488, 517–521 (2012).

16. Halbleib, J.M. & Nelson, W.J. Cadherins in development: cell adhesion, sorting, and tissue morphogenesis. Genes & development 20, 3199–3214 (2006).

17. Thomson, J.A. et al. Embryonic stem cell lines derived from human blastocysts. Science 282, 1145–1147 (1998).

18. Clevers, H. Modeling Development and Disease with Organoids. Cell 165, 1586–1597 (2016).

19. West, M.D. et al. Use of deep neural network ensembles to identify embryonic-fetal transition markers: repression of COX7A1 in embryonic and cancer cells. Oncotarget 9, 7796–7811 (2018).

20. West, M.D. et al. The ACTCellerate initiative: large-scale combinatorial cloning of novel human embryonic stem cell derivatives. Regen Med 3, 287–308 (2008).

21. Buenrostro, J.D., Giresi, P.G., Zaba, L.C., Chang, H.Y. & Greenleaf, W.J. Transposition of native chromatin for fast and sensitive epigenomic profiling of open chromatin, DNA-binding proteins and nucleosome position. Nat Methods 10, 1213–1218 (2013).

22. Yang, J. & Corces, V.G. Insulators, long-range interactions, and genome function. Curr Opin Genet Dev 22, 86–92 (2012).

23. Sternberg, H. et al. A human embryonic stem cell-derived clonal progenitor cell line with chondrogenic potential and markers of craniofacial mesenchyme. Regen Med 7, 481–501 (2012).

24. West, M.D. et al. Clonal derivation of white and brown adipocyte progenitor cell lines from human pluripotent stem cells. Stem cell research & therapy 10, 7 (2019).

25. Toyota, M. et al. CpG island methylator phenotype in colorectal cancer. Proc Natl Acad Sci U S A 96, 8681–8686 (1999).

26. Ogino, S. et al. Evaluation of markers for CpG island methylator phenotype (CIMP) in colorectal cancer by a large population-based sample. J Mol Diagn 9, 305–314 (2007).

27. Lund, E.G., Duband-Goulet, I., Oldenburg, A., Buendia, B. & Collas, P. Distinct features of lamin A-interacting chromatin domains mapped by ChIP-sequencing from sonicated or micrococcal nuclease-digested chromatin. Nucleus 6, 30–39 (2015).

28. Guelen, L. et al. Domain organization of human chromosomes revealed by mapping of nuclear lamina interactions. Nature 453, 948–951 (2008).

29. Khan, A. & Zhang, X. dbSUPER: a database of super-enhancers in mouse and human genome. Nucleic Acids Res 44, D164–171 (2016).

30. Kim, S., Yu, N.K. & Kaang, B.K. CTCF as a multifunctional protein in genome regulation and gene expression. Exp Mol Med 47, e166 (2015).

31. Igolkina, A.A. et al. H3K4me3, H3K9ac, H3K27ac, H3K27me3 and H3K9me3 Histone Tags Suggest Distinct Regulatory Evolution of Open and Condensed Chromatin Landmarks. Cells 8 (2019).

32. Goodman, K.M. et al. Protocadherin cis-dimer architecture and recognition unit diversity. Proc Natl Acad Sci U S A 114, E9829–E9837 (2017).

33. Kehayova, P., Monahan, K., Chen, W. & Maniatis, T. Regulatory elements required for the activation and repression of the protocadherin-alpha gene cluster. Proc Natl Acad Sci U S A 108, 17195–17200 (2011).

34. Sternberg, H. et al. Seven diverse human embryonic stem cell-derived chondrogenic clonal embryonic progenitor cell lines display site-specific cell fates. Regen Med (2012).

35. Jakab, K. et al. Tissue engineering by self-assembly and bio-printing of living cells. Biofabrication 2, 022001 (2010).

36. Constantinescu, D., Gray, H.L., Sammak, P.J., Schatten, G.P. & Csoka, A.B. Lamin A/C expression is a marker of mouse and human embryonic stem cell differentiation. Stem Cells 24, 177–185 (2006).

37. Shimi, T. et al. The role of nuclear lamin B1 in cell proliferation and senescence. Genes & development 25, 2579–2593 (2011).

38. West, M.D. et al. Toward a unified theory of aging and regeneration. Regen Med 14, 867–886 (2019).

39. Sternberg, H., Janus, J. & West, M.D. Defining cell-matrix combination products in the era of pluripotency. Biomatter 3 (2013).

40. Almenar-Queralt, A. et al. Chromatin establishes an immature version of neuronal protocadherin selection during the naive-to-primed conversion of pluripotent stem cells. Nat Genet 51, 1691–1701 (2019).

41. Kaczkowski, B. et al. Transcriptome Analysis of Recurrently Deregulated Genes across Multiple Cancers Identifies New Pan-Cancer Biomarkers. Cancer research 76, 216–226 (2016).

42. K., V.R.L. Cellular Pathology (Originally Published 1859). (Classics of Medicine, U.S.A.; 1978).

43. Warburg, O. On the origin of cancer cells. Science 123, 309–314 (1956).

44. Manzo, G. Similarities Between Embryo Development and Cancer Process Suggest New Strategies for Research and Therapy of Tumors: A New Point of View. Front Cell Dev Biol 7, 20 (2019).

45. Abe, M. et al. CpG island methylator phenotype is a strong determinant of poor prognosis in neuroblastomas. Cancer research 65, 828–834 (2005).

46. Henrich, K.O. et al. Integrative Genome-Scale Analysis Identifies Epigenetic Mechanisms of Transcriptional Deregulation in Unfavorable Neuroblastomas. Cancer research 76, 5523–5537 (2016).

47. Vega-Benedetti, A.F. et al. Clustered protocadherins methylation alterations in cancer. Clin Epigenetics 11, 100 (2019).

48. West, M.D. et al. The germline/soma dichotomy: implications for aging and degenerative disease. Regen Med 11, 331–334 (2016).

49. Rober, R.A., Weber, K. & Osborn, M. Differential timing of nuclear lamin A/C expression in the various organs of the mouse embryo and the young animal: a developmental study. Development 105, 365–378 (1989).

50. Solovei, I. et al. LBR and lamin A/C sequentially tether peripheral heterochromatin and inversely regulate differentiation. Cell 152, 584–598 (2013).

51. Tardat, M. & Dejardin, J. Telomere chromatin establishment and its maintenance during mammalian development. Chromosoma 127, 3–18 (2018).

52. van Steensel, B. & Belmont, A.S. Lamina-Associated Domains: Links with Chromosome Architecture, Heterochromatin, and Gene Repression. Cell 169, 780–791 (2017).

53. Salpea, P. et al. Postnatal development- and age-related changes in DNA-methylation patterns in the human genome. Nucleic Acids Res 40, 6477–6494 (2012).

54. Matsuyama, M. et al. Analysis of epigenetic aging in vivo and in vitro: Factors controlling the speed and direction. Exp Biol Med (Maywood) 245, 1543–1551 (2020).

55. Dixon, J.R. et al. Chromatin architecture reorganization during stem cell differentiation. Nature 518, 331–336 (2015).

56. Mah, K.M., Houston, D.W. & Weiner, J.A. The gamma-Protocadherin-C3 isoform inhibits canonical Wnt signalling by binding to and stabilizing Axin1 at the membrane. Sci Rep 6, 31665 (2016).

57. Ocampo, A. et al. In Vivo Amelioration of Age-Associated Hallmarks by Partial Reprogramming. Cell 167, 1719–1733 e1712 (2016).

58. Lanza, R.P., Cibelli, J.B. & West, M.D. Human therapeutic cloning. Nat Med 5, 975–977 (1999).

59. Plath, K. & Lowry, W.E. Progress in understanding reprogramming to the induced pluripotent state. Nat Rev Genet 12, 253–265 (2011).

60. Soufi, A., Donahue, G. & Zaret, K.S. Facilitators and impediments of the pluripotency reprogramming factors’ initial engagement with the genome. Cell 151, 994–1004 (2012).

61. Trapnell, C. et al. Differential gene and transcript expression analysis of RNA-seq experiments with TopHat and Cufflinks. Nat Protoc 7, 562–578 (2012).

62. Corces, M.R. et al. An improved ATAC-seq protocol reduces background and enables interrogation of frozen tissues. Nat Methods 14, 959–962 (2017).

63. Bentsen, M. et al. ATAC-seq footprinting unravels kinetics of transcription factor binding during zygotic genome activation. Nat Commun 11, 4267 (2020).

64. Zhang, Y. et al. Model-based analysis of ChIP-Seq (MACS). Genome Biol 9, R137 (2008).

65. Fornes, O., et al. JASPAR 2020: update of the open-access database of transcription factor binding profiles. Nucleic Acids Res 48, D87–D92 (2020).

66. Robinson, J.T., et al. Integrative genomics viewer. Nat Biotechnol 29, 24–26 (2011).

67. Krueger, F. & Andrews, S.R. Bismark: a flexible aligner and methylation caller for Bisulfite-Seq applications. Bioinformatics 27, 1571–1572 (2011).

68. Juhling, F. et al. metilene: fast and sensitive calling of differentially methylated regions from bisulfite sequencing data. Genome Res 26, 256–262 (2016).

69. Lieberman-Aiden, E. et al. Comprehensive mapping of long-range interactions reveals folding principles of the human genome. Science 326, 289–293 (2009).

70. Li, H. & Durbin, R. Fast and accurate long-read alignment with Burrows-Wheeler transform. Bioinformatics 26, 589–595 (2010).

71. Faust, G.G. & Hall, I.M. SAMBLASTER: fast duplicate marking and structural variant read extraction. Bioinformatics 30, 2503–2505 (2014).

72. Li, H. et al. The Sequence Alignment/Map format and SAMtools. Bioinformatics 25, 2078–2079 (2009).

73. Shin, H. et al. TopDom: an efficient and deterministic method for identifying topological domains in genomes. Nucleic Acids Res 44, e70 (2016).

74. J.D., H. Matplotlib: a 2D graphics environment. Computing in Science & Engineering 9, 90–95 (2007).

75. Durand, N.C. et al. Juicebox Provides a Visualization System for Hi-C Contact Maps with Unlimited Zoom. Cell Syst 3, 99–101 (2016).

76. Zang et al. A clustering approach for identification of enriched domains from histone modification ChIP-Seq data. Bioinformatics 25, 1952–1958 (2009).

