## Supplementary material for "Differential Expression of α, β, and γ Protocadherin Isoforms During Differentiation, Aging, and Cancer": Differential Expression of Clustered PCDH-Supplementary Materials

### Supplementary Figure 1A-C

## A.

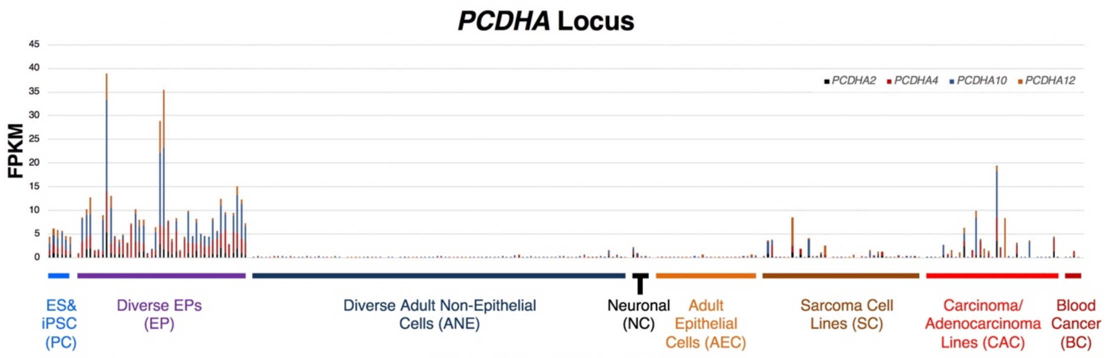


## B.

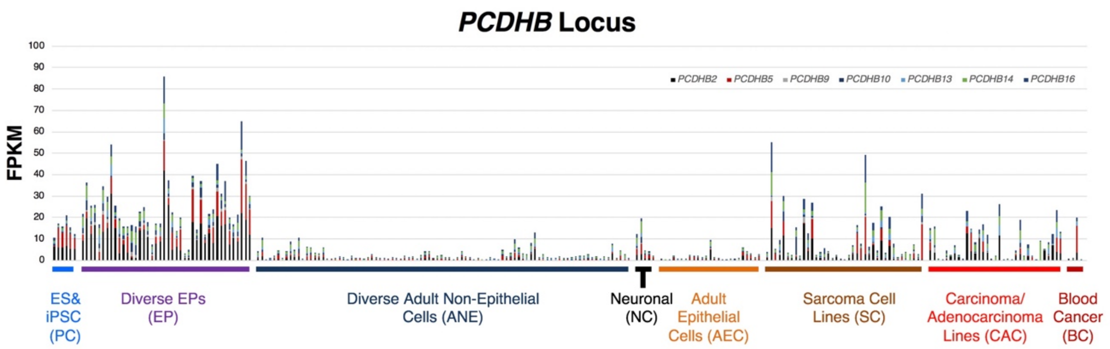


## C.

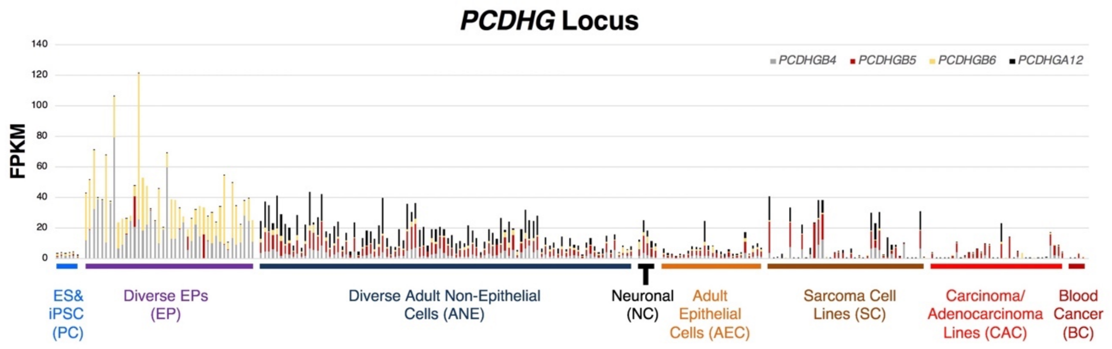


**Supplementary Figure 1. CPL isoform expression in Pluripotent Cells (PC), Embryonic Progenitors (EP), Adult Non-Epithelial (ANE), Neuronal Cells (NC), Adult Epithelial Cells**

**(AEC), Sarcoma Cells (SC), Carcinoma and Adenocarcinoma Cells (CAC), and Blood Cell (BC) cancer lines.** A) Cumulative Expression of statistically-significantly differentially-expressed α cluster genes: *PCDHA2*, *PCDHA4*, *PCDHA10*, and *PCDHA12*. B). Analogous cumulative expression of the β cluster genes: *PCDHB2*, *PCDHB5*, *PCDHB9*, *PCDHB10, PCDHB13*, *PCDHB14*, and *PCDHB16.* C). Analogous cumulative expression of the γ cluster genes: *PCDHGB4*, *PCDHGB5*, *PCDHGB6*, and *PCDHGA12.*

#### Supplementary Figure 2

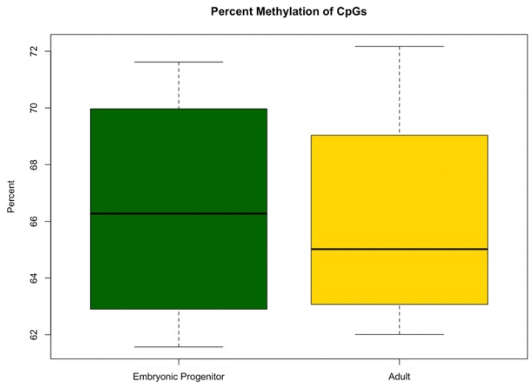


**Supplementary Figure 2. Percent global CpG methylation in embryonic progenitors vs adult counterparts.** The percent of all CpG dinucleotides that are methylated in four hES cell-derived clonal embryonic progenitor (EP) cell lines was tabulated together with their four respective adult-derived counterparts: 4D20.8, a clonal embryonic osteochondral progenitor and adult bone marrow-derived mesenchymal stem cells (MSCs); 30-MV2-6, a clonal embryonic vascular endothelial cell line together with adult-derived aortic endothelial cells (HAECs); SK5 a clonal embryonic skeletal muscle progenitor together with adult skeletal myoblasts; and E3, a clonal embryonic white preadipocyte cell line together with adult subcutaneous white preadipocytes. (Error bars represent standard error of the mean).

### Supplementary Figure 3

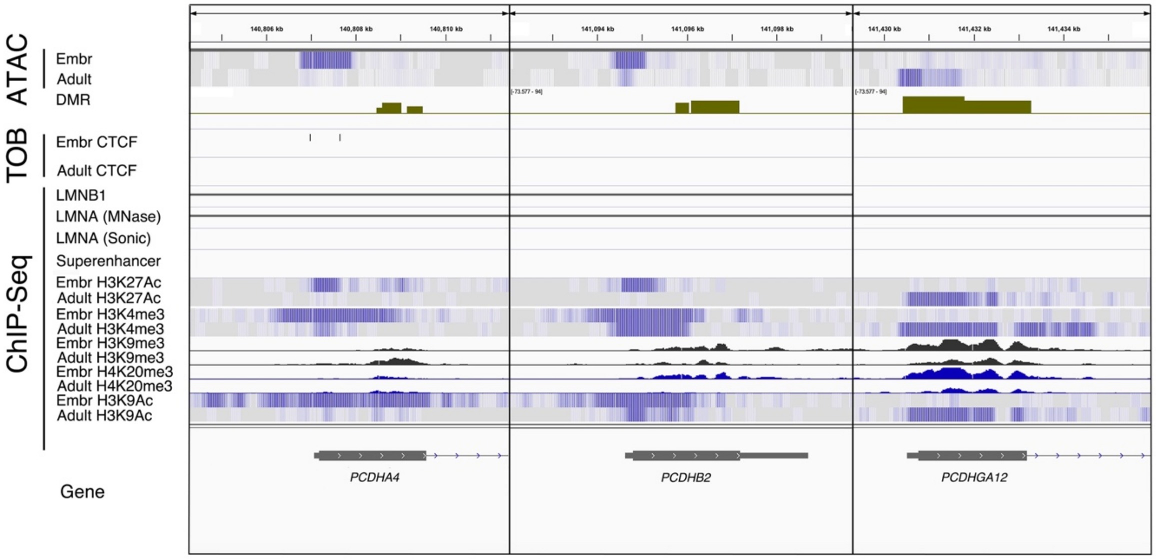


**Supplementary Figure 3. Comparison of epigenetic markers in the 5' region of *PCDHA4, PCDHB2*, and *PCDHGA12*.** Rows represent: 1) ATAC-seq of the embryonic progenitor osteogenic mesenchymal line 4D20.8 (Embr Mesen) and adult-derived osteogenic mesenchyme (MSCs). 2) Differentially Methylated Regions (DMRs) are shown where elevated green represents hypermethylation in embryonic cells and depressed red signal represents relative hypomethylation in embryonic compared to adult cells; 3) ChIP-seq values for LMNB1, LMNA (micrococcal nuclease treated (MNase), LMNA (sonicated); 4) ChIP-seq data in paired embryonic and adult mesenchymal cells resulting from precipitation of chromatin with antibodies directed to H3K27Ac, H2K4me3, H2K9me3, and H4K20me3, and H3K9Ac for the entire CPL. (hg38 Chromosome position: Chr5: 140,650,000-141,700,000), image captured from IGV.

### Supplementary Figure 4A-B

## A.

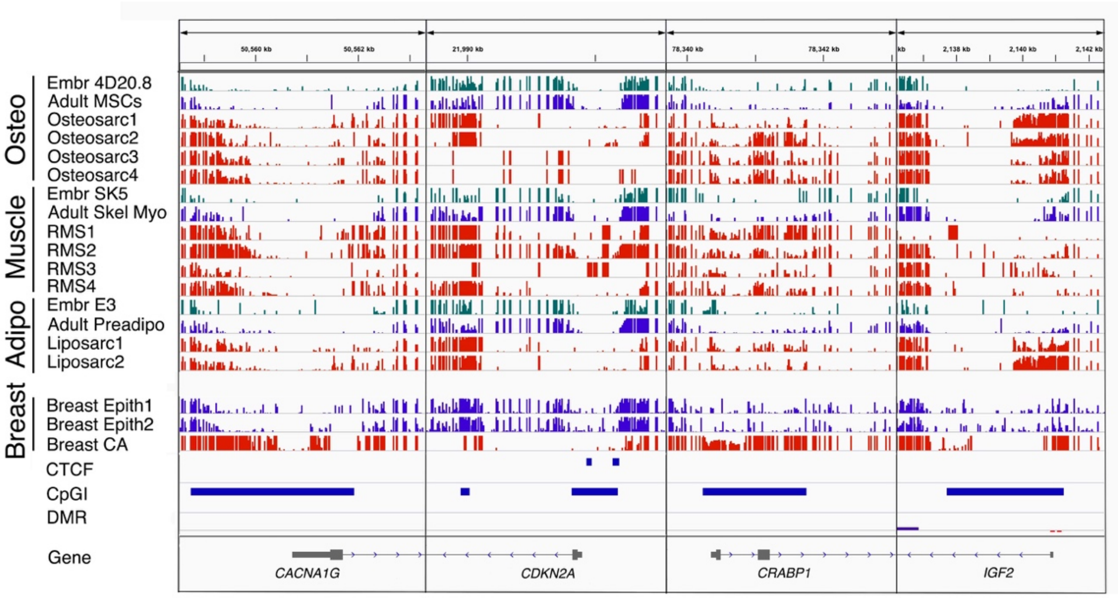


## B.

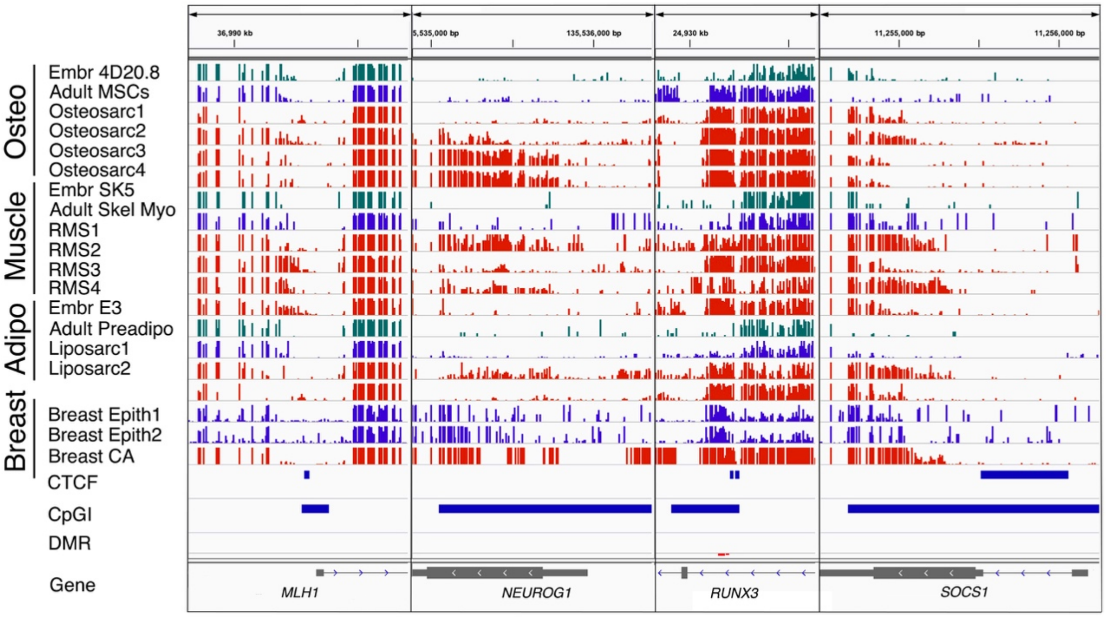


**Supplementary Figure 4. WGBS results for CIMP loci in select EP, ANE, and SC, and CAC cell lines.** 1) WGBS results showing percent methylation of CpGs in three EP lines (shown in green): Adult osteogenic mesenchyme 4D20.8 (Embr Mesen); Embryonic skeletal myoblasts. (SK5); and Embryonic white preadipocytes (E3); Adult counterparts (shown in blue) being: Adult osteogenic mesenchyme (MSCs); Adult skeletal myoblasts (Adult skel myo); and Adult white preadipocytes (Adult preadipo); and sarcoma counterparts (shown in red) including four osteosarcoma cell lines (Osteosarc1-4), four rhabdomyosarcoma lines (RMS1-4), two liposarcoma lines (Liposarc1-2), also displayed are two normal adult breast epithelial cell lines (shown in blue) and MCF7 breast cancer cells (shown in red). 2) CTCF binding sites, 3) CpG islands (CpGIs), 4) Differentially Methylated Regions (DMRs) are shown where elevated green represents hypermethylation in embryonic cells and depressed red signal represents relative hypomethylation in embryonic compared to adult cells. A) Shown are 5' regions for the CIMP genes: *CACNA1G, CDKN2A, CRABP1*, and *IGF2*. B) Shown are 5' regions for the CIMP genes: *MLH1, NEUROG1, RUNX3*, and *SOCS1*. (hg38 Chromosome position: Chr5: 140,650,000-141,700,000), image captured from IGV.

#### Supplementary Figure 5A-B

**A.**

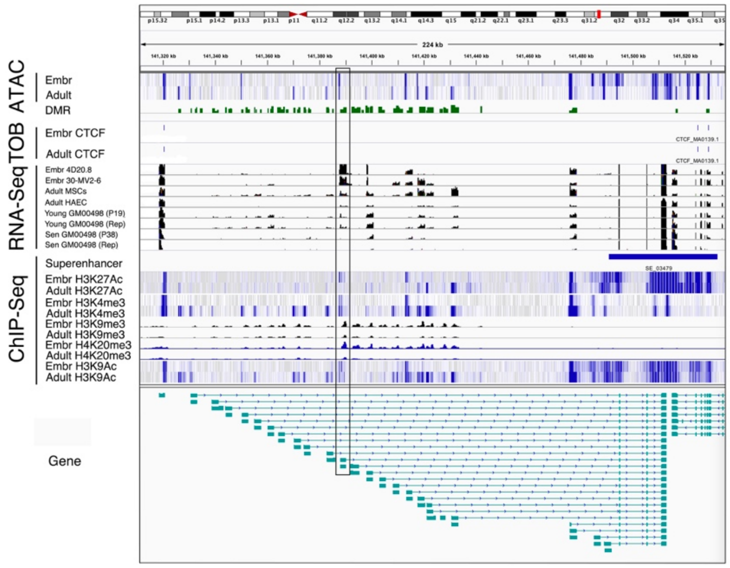


**B.**

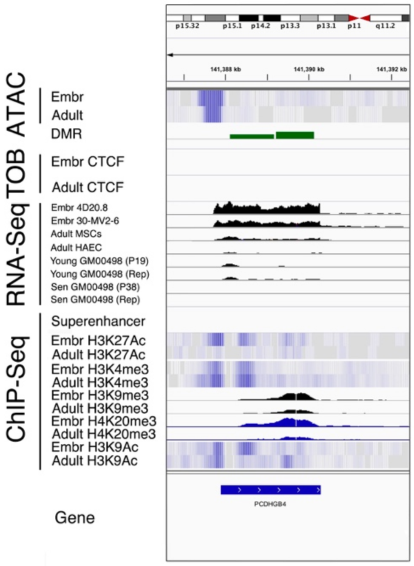


**Supplementary Figure 5. RNA-Sequence data from the γ cluster during development and aging *in vitro*.** Rows represent: 1) ATAC-seq of the embryonic progenitor osteogenic mesenchymal line 4D20.8 (Embr) and adult-derived osteogenic mesenchyme (MSCs) (Adult). 2) Differentially Methylated Regions (DMRs) are shown where elevated green represents hypermethylation in embryonic cells and depressed red signal represents relative hypomethylation in embryonic compared to adult cells; 3) CTCF binding sites in Embr Mesen and Adult Mesen based on TOBIAS analysis of ATAC footprints; 4) RNA-sequence coverage in embryonic progenitors to osteogenic mesenchyme and vascular endothelium (Embr 4D20.8 and Embr 30-MV2-6 respectively) and adult-derived osteogenic mesenchyme and aortic endothelial cells (Adult MSCs and Adult HAEC respectively); Young (P19) GM00498 fibroblasts and a biological replicate and senescent (P38) GM00498 fibroblasts and a biological replicate; 5) location of superenhancers in the γ region ChIP-seq data in paired embryonic and adult cells resulting from precipitation of chromatin with antibodies directed to H3K27Ac, H2K4me3,

H2K9me3, and H4K20me3, and H3K9Ac for the γ locus. A) Entire γ cluster. B) Zoom in on region on first exon of *PCDHGB4*. (hg38 genome, image captured from IGV.

### Supplementary Figure 6A-B

**A.**

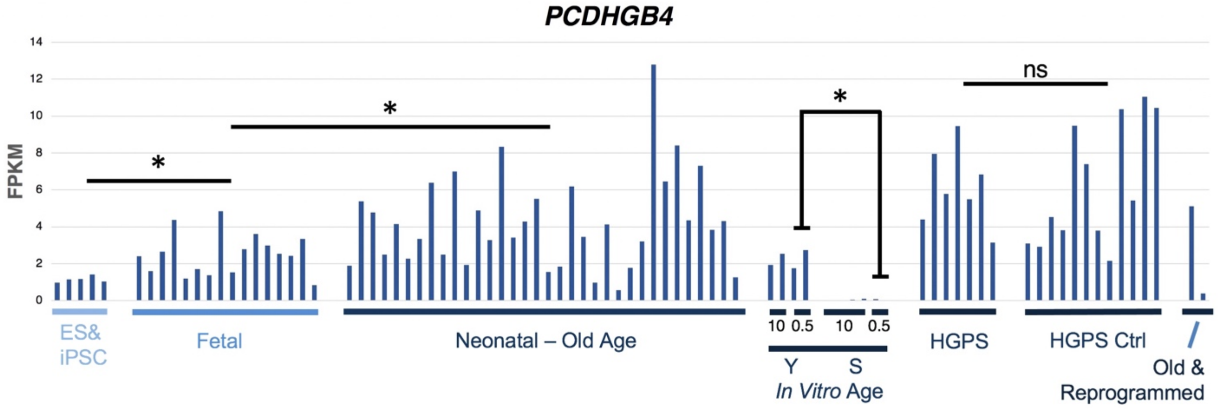


**B.**

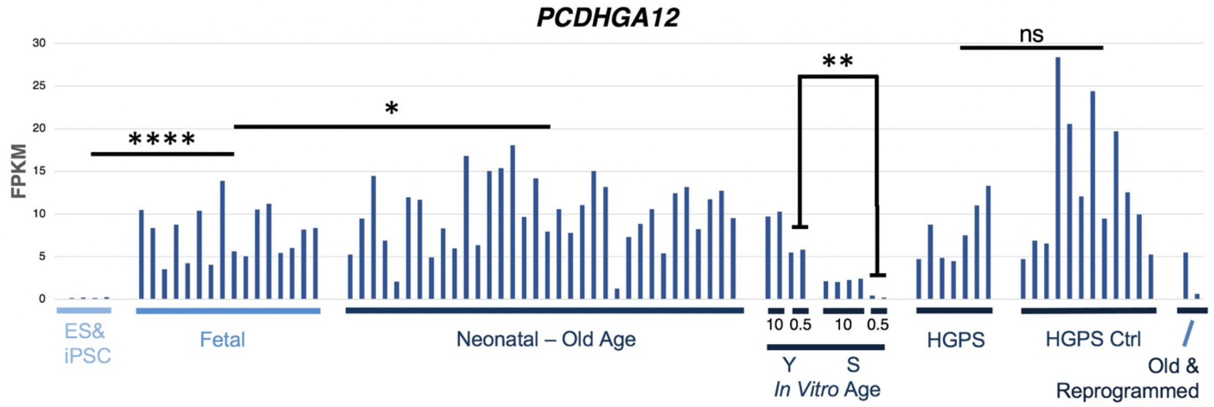


**Supplementary Figure 6. RNA-Sequence data for *PCDHGB4* and *PCDHGA12* expression during fibroblast aging *in vivo* and *in vitro*.** A) Expression of *PCDHGB4* is shown in pluripotent stem cells including hES and hiPS cells; fetal-derived early passage dermal fibroblasts spanning 8-16 weeks of gestation synchronized for five days in quiescence; adult-derived dermal fibroblasts spanning 11-83 years of age synchronized five days in quiescence; *In vitro* aged GM00498 dermal fibroblasts from a 3-year old donor at early passage (P19) harvested in log growth (10% FBS) or quiescence induced by culture in 0.5% FBS for two weeks and at late passage (P38) harvested in log growth or quiescence for two weeks; dermal fibroblasts from Hutchinson-Gilford Progeria Syndrome (HGPS) patients vs normal age-matched controls synchronized five days in quiescence; and dermal fibroblasts from a 59 year-old donor and same cells transcriptionally reprogramed in pluripotency (iPS cells). (ns: not significant) (* $p<0.05$), (** $p<0.01$), (**** $p< 0.0001$).

### Supplementary Figure 7A-B

## A.

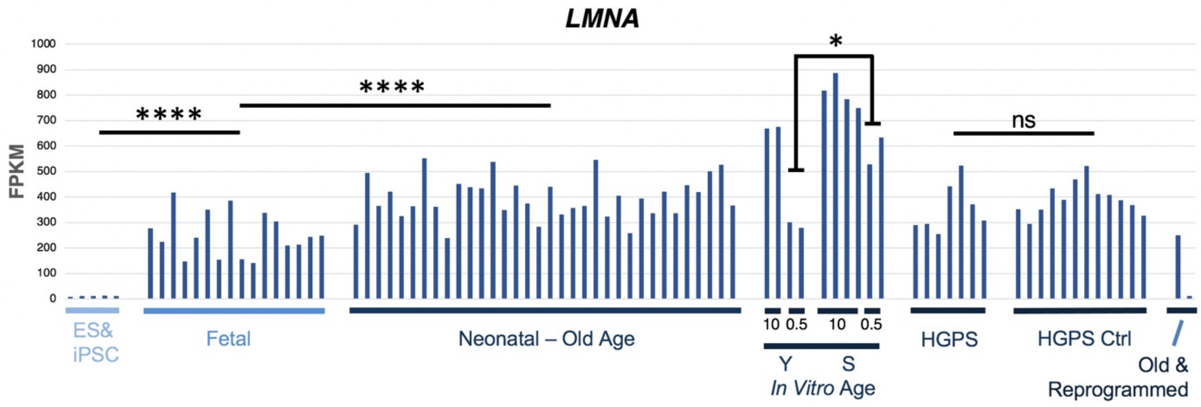


## B.

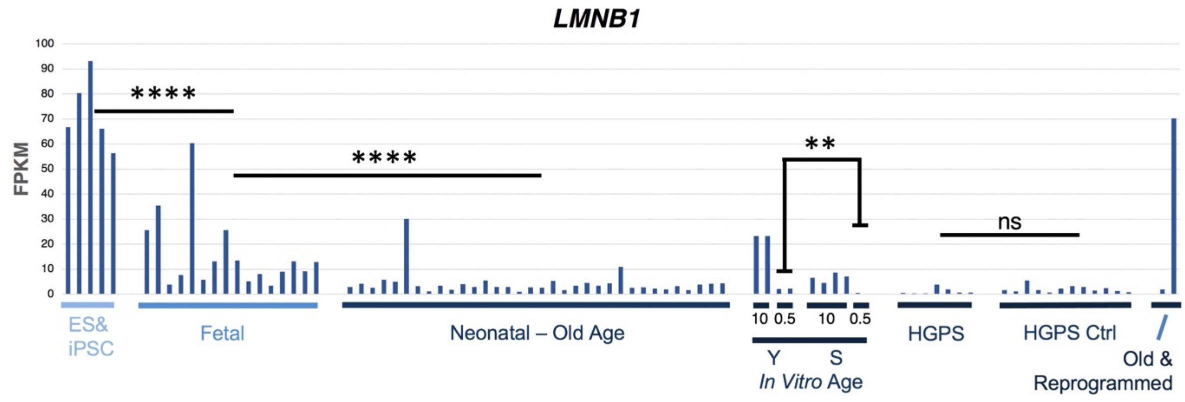


**Supplementary Figure 7. RNA-Sequence data for *LMNA* and *LMNB1* expression during fibroblast aging *in vivo* and *in vitro*.** A) Expression of *LMNA* is shown in pluripotent stem cells including hES and hiPS cells; fetal-derived early passage dermal fibroblasts spanning 8-16 weeks of gestation synchronized for five days in quiescence; adult-derived dermal fibroblasts spanning 11-83 years of age synchronized five days in quiescence; *In vitro* aged GM00498 dermal fibroblasts from a 3-year old donor at early passage (P19) harvested in log growth (10% FBS) or quiescence induced by culture in 0.5% FBS for two weeks and at late passage (P38)

harvested in log growth or quiescence for two weeks; dermal fibroblasts from Hutchinson-Gilford Progeria Syndrome (HGPS) patients vs normal age-matched controls synchronized five days in quiescence; and dermal fibroblasts from a 59 year-old donor and same cells transcriptionally reprogramed in pluripotency (iPS cells). (ns: not significant) (* $p<0.05$), (** $p<0.01$), (**** $p< 0.0001$).

### Supplementary Figure 8A-B

## A.

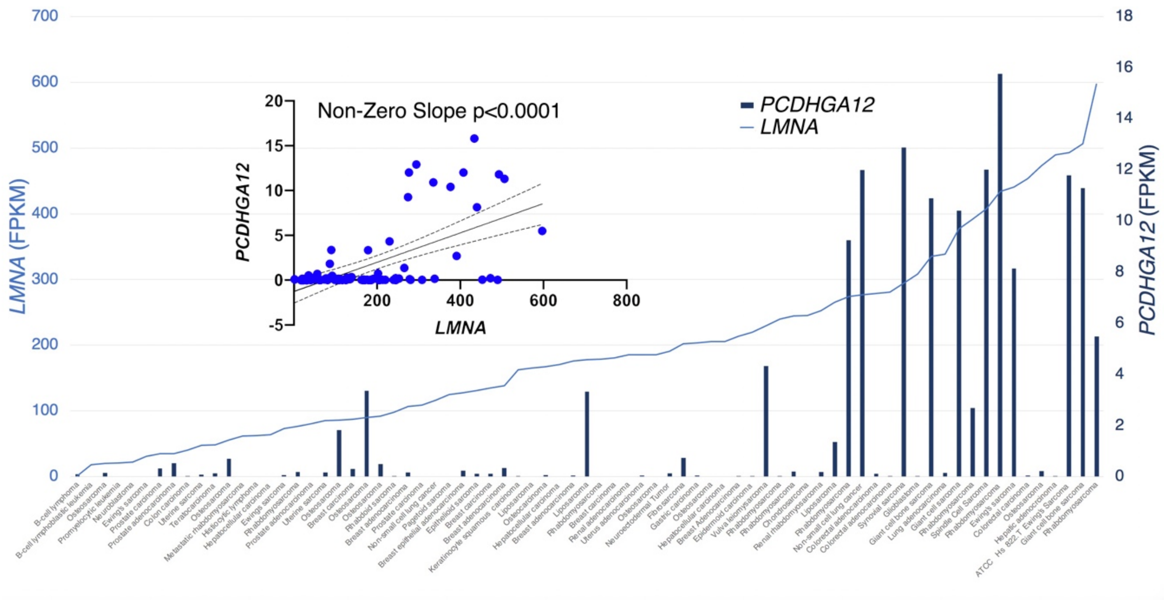


## B.

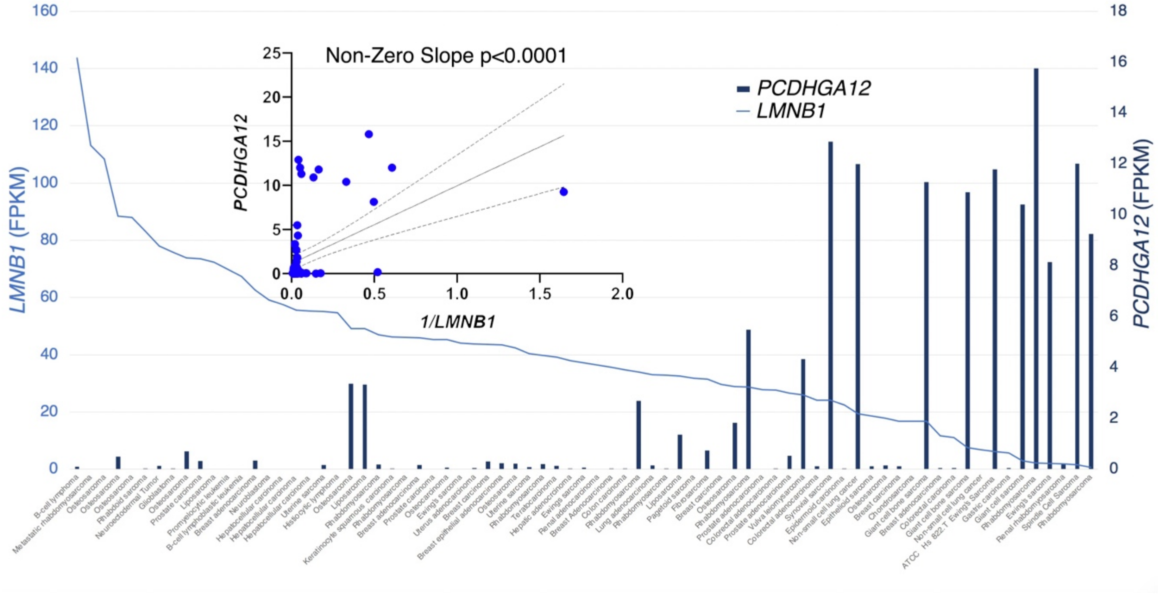


**Supplementary Figure 8. Correlation of *PCDHGA12* with LMNA and LMNB1 expression in cancer cell lines.** A) Expression in FPKM of *PCDHGA12* in diverse sarcoma and carcinoma cell lines sorted by *LMNA* expression. (Insert, linear regression with positive correlation (p<0.0001). B) Expression in FPKM of *PCDHGA12* in diverse sarcoma and carcinoma cell lines sorted by *LMNB1* expression. (Insert, linear regression with inverse correlation (p<0.0001).

**Supplementary Table I. Description and origin of cell lines.**

| Cell Type | Cell Line Designation | Source |
|---|---|---|
| hES cells (PCs) | H9 P41 matrigel | Wisconsin Alumni Research Foundation |
| | MA03 P18 matrigel | Astellas Istitute for Renerative Medicine |
| | ESI 017 P24 matrigel | ESI-Lineage Cell Therapeutics |
| | ESI 053 P23 matrigel | ESI-Lineage Cell Therapeutics |
| Human embryonic progenitor-derived iPS cells (PCs) | EH3 P4 iPS-H9 matrigel | Lineage Cell Therapeutics Inc |
| Adult dermal fibroblast cells | MDW-1 P7 control | Lineage Cell Therapeutics Inc |
| Adult dermal fibroblast cell-derived iPS Cells (PCs) | MDW-1 iPS cl.1 P6 | Lineage Cell Therapeutics Inc |
| Diverse human clonal Embryonic Progenitor (EP) cell lines | EN13 P14 ctrl | Lineage Cell Therapeutics Inc |
| | 7SMOO32 P11 ctrl | Lineage Cell Therapeutics Inc |
| | SM30 P15 ctrl | Lineage Cell Therapeutics Inc |
| | 4D20.8 P15 ctrl | Lineage Cell Therapeutics Inc |
| | E15 P19 ctrl | Lineage Cell Therapeutics Inc |
| | 7PEND24 P21 ctrl | Lineage Cell Therapeutics Inc |
| | SK11 P16 ctrl | Lineage Cell Therapeutics Inc |
| | MEL2 P17 ctrl | Lineage Cell Therapeutics Inc |
| | T42 P17 ctrl | Lineage Cell Therapeutics Inc |
| | E69 P16 ctrl | Lineage Cell Therapeutics Inc |
| | E3 P12 ctrl | Lineage Cell Therapeutics Inc |
| | C4ELS5.1 P14 ctrl | Lineage Cell Therapeutics Inc |
| | ESI004 NP110 SM P12 ctrl | Lineage Cell Therapeutics Inc |
| | ESI004 NP88 SM P12 ctrl | Lineage Cell Therapeutics Inc |
| | 7SMOO32 P13 D14 MM | Lineage Cell Therapeutics Inc |
| | SM30 P14 D14 Chondro MM | Lineage Cell Therapeutics Inc |
| | 4D20.8 P20 D14 Hystem TGFb3 10ng/ml + BMP4 10ng/ml | Lineage Cell Therapeutics Inc |
| | E15 P22 D14 Chondro Hystem BMP4 10ng/ml + TGFb3 10ng/ml | Lineage Cell Therapeutics Inc |
| | 7PEND24 P22 D14 Hystem BMP4 10ng/ml + TGFb3 10ng/ml | Lineage Cell Therapeutics Inc |
| | SK11 P16 D14 Hystem BMP4 10ng/ml + TGFb3 10ng/ml | Lineage Cell Therapeutics Inc |
| | MEL2 P26 D14 Hystem BMP2 50ng/ml + TGFb3 10ng/ml | Lineage Cell Therapeutics Inc |
| | T42 P17 D14 MM BMP4 10ng/ml | Lineage Cell Therapeutics Inc |
| | E69 P16 D14 MM BMP4 10ng/ml | Lineage Cell Therapeutics Inc |
| | E3 P17 D21 confluence BMP4 50ng/ml + Rosiglitazone 5uM | Lineage Cell Therapeutics Inc |
| | C4ELS5.1 P14 D14 Hystem rosiglitazone 1uM + T3 2nM + last 4 hours CL316243 10uM | Lineage Cell Therapeutics Inc |
| | ESI004 NP110 SM P17 D14 Hystem rosiglitazone 1uM + T3 2nM + last 4 hours CL316243 10uM | Lineage Cell Therapeutics Inc |
| | ESI004 NP88 SM P15 D14 Hystem rosiglitazone 1uM + T3 2nM + last 4 hours CL316243 10uM | Lineage Cell Therapeutics Inc |
| | 30-MV2-6 P6 | Lineage Cell Therapeutics Inc |
| | 30-MV2-3 P6 | Lineage Cell Therapeutics Inc |
| | 30-MV2-4 P6 | Lineage Cell Therapeutics Inc |
| | 30-MV2-10 P6 | Lineage Cell Therapeutics Inc |
| | 30-MV2-17 P6 | Lineage Cell Therapeutics Inc |

| | | |
|---|---|---|
| | 30-MV2-19 P6 | Lineage Cell Therapeutics Inc |
| | 30-MV2-2 P6 | Lineage Cell Therapeutics Inc |
| | 30-MV2-7 P6 | Lineage Cell Therapeutics Inc |
| | 30-MV2-9 P6 | Lineage Cell Therapeutics Inc |
| | 30-MV2-14 P6 | Lineage Cell Therapeutics Inc |
| | 30-MV2-24 P6 | Lineage Cell Therapeutics Inc |
| | 30-MV2-8 P6 | Lineage Cell Therapeutics Inc |
| | 30-SM2-1 P6 | Lineage Cell Therapeutics Inc |
| | 30-SM2-3 P6 | Lineage Cell Therapeutics Inc |
| | RP1-SKEL-8 P6 | Lineage Cell Therapeutics Inc |
| **Fetal cell lines (FCs)** | Fetal brown preadipocytes (Zenbio) P4 ctrl | ZenBio |
| | Human fetal brown adipocytes 20 wk Zenbio RNA-T10-CS (Lot 10914A) | ZenBio |
| | Fetal brown preadipocytes (Zenbio) P8 ctrl | ZenBio |
| | Fetal dermal fibroblasts P3 arm FB 8wk | ZenBio |
| | Fetal dermal fibroblasts P4 arm FB 9wk | ZenBio |
| | Fetal dermal fibroblasts P2 arm FB 10wk | ZenBio |
| | Fetal dermal fibroblasts P2 arm FB 11wk | ZenBio |
| | Fetal dermal fibroblasts P2 arm FB 16wk | ZenBio |
| | Neonatal foreskin fibroblasts (Xgene) P17 ctrl | X-Gene |
| | Normal human articular chondrocytes P6 | Lonza |
| | Human umbilical artery smooth muscle cells P5 | ScienCell Research Laboratories |
| | Coronary artery smooth muscle cells P14 | Lonza |
| | HSC human schwann cell P2 | ScienCell Research Laboratories |
| | Brain Pericytes P6 | ScienCell Research Laboratories |
| | Human brain vascular smooth muscle cells P3 | ScienCell Research Laboratories |
| | Skeletal myoblasts Zenbio P5 ctrl | ZenBio |
| | Omental Preadipocyte Zenbio (Lot SLOM-18) P9 ctrl | ZenBio |
| | Subcutaneous preadipocytes Zenbio (Lot SLOO54) P6 ctrl | ZenBio |
| | HSMM (Lonza human skeletal muscle myoblasts) P4 ctrl | Lonza |
| | NHEM (Lonza neonatal human epidermal melanocytes) P7 ctrl | Lonza |
| | NHOST Lonza normal human osteoblasts) P5 ctrl | Lonza |
| | Dental pulp stem cells P6 ctrl | Songtau Shi USC School of Dentistry |
| | HAoSMC (aortic smooth muscle PromoCell) lot 4002012.2 P5 ctrl | PromoCell |
| | HHORSC t Human Hair Outer Root Sheath Cell, MBA_2123 | ScienCell Research Laboratories |
| | HTMC t Human Trabecular Meshwork Cell, MBA_2174 | ScienCell Research Laboratories |
| | HSC t Human Schwann Cell, MBA_2107 | ScienCell Research Laboratories |
| | HBVSMC t Human Brain Vascular Smooth Muscle Cell, MBA_2101 | ScienCell Research Laboratories |
| | HBVP t Human Brain Vascular Pericyte, MBA_2102 | ScienCell Research Laboratories |
| | HCPF t Human Choroid Plexus Fibroblast, MBA_2105 | ScienCell Research Laboratories |
| | HHDPC t Human Hair Dermal Papilla Cell 10µg, MBA_2122 | ScienCell Research Laboratories |
| | HLF t Human Lymphatic Fibroblasts Cell, MBA_2125 | ScienCell Research Laboratories |

| | | |
|---|---|---|
| Diverse Adult Non-Epithelial (stromal & parenchymal) cell types (ANEs) | HPLF t Human Peridontal Ligament Fibroblast, MBA_2127 | ScienCell Research Laboratories |
| | HISMC t Human Intestinal Smooth Muscle Cell, MBA_2130 | ScienCell Research Laboratories |
| | HCoSMC t Human Colonic Smooth Muscle Cell, MBA_2131 | ScienCell Research Laboratories |
| | HPASMC t Human Pulmonary Artery Smooth Muscle Cell, MBA_2134 | ScienCell Research Laboratories |
| | HPAF t Human Pulmonary Artery Fibroblast, MBA_2135 | ScienCell Research Laboratories |
| | HPF t Human Pulmonary Fibroblast, MBA_2140 | ScienCell Research Laboratories |
| | HBSM t Human Bronchial Smooth Muscle Cell, MBA_2141 | ScienCell Research Laboratories |
| | HTSMC t Human Tracheal Smooth Muscle Cell, MBA_2142 | ScienCell Research Laboratories |
| | HSkMC t Human Skeletal Muscle Cell, MBA_2143 | ScienCell Research Laboratories |
| | HSkMM t Human Skeletal Muscle Myoblast, MBA_2145 | ScienCell Research Laboratories |
| | HRMC t Human Renal Mesangial Cell, MBA_2149 | ScienCell Research Laboratories |
| | HBdSMC t Human Bladder Smooth Muscle Cell, MBA_2151 | ScienCell Research Laboratories |
| | HPrF tDNA Human Prostate Fibroblasts Cell, MBA_2154 | ScienCell Research Laboratories |
| | HCO t Human Calvarial Osteoblasts Cell, MBA_2155 | ScienCell Research Laboratories |
| | HNPC tDNA Human Nucleus Pulposus Cells, MBA_2157 | ScienCell Research Laboratories |
| | HAFC t Human Annulus Fibrosus Cell, MBA_2158 | ScienCell Research Laboratories |
| | HHSteC t Human Hepatic Stellate Cell, MBA_2161 | ScienCell Research Laboratories |
| | HCM-a t Human Cardiac Myocyte-adult, MBA_2165 | ScienCell Research Laboratories |
| | HCF-av t Human Cardiac Fibroblast-adult ventrical, MBA_2166 | ScienCell Research Laboratories |
| | HConF t Human Conjunctival Fibroblast, MBA_2172 | ScienCell Research Laboratories |
| | HVMF t Human Villous Mesenchymal Fibroblast, MBA_2176 | ScienCell Research Laboratories |
| | HAmMSC t Human Amniotic Mesenchymal Stromal Cell, MBA_2177 | ScienCell Research Laboratories |
| | HCMSC t Human Chorionic Mesenchymal Stromal Cell, MBA_2178 | ScienCell Research Laboratories |
| | HPA-v t Human Preadipocyte-visceral, MBA_2179 | ScienCell Research Laboratories |
| | HPA-s t Human Preadipocyte-subcutaneous, MBA_2180 | ScienCell Research Laboratories |
| | HOF t Human Ovarian Fibroblast t, MBA_2182 | ScienCell Research Laboratories |
| | HMSC-bm t Human Mesenchymal Stem Cell-bone marrow, MBA_2183 | ScienCell Research Laboratories |
| | HMSC-ad t Human Mesenchymal Stem Cell-adipose, MBA_2184 | ScienCell Research Laboratories |
| | HMSC-he t Human Mesenchymal Stem Cell-hepatic, MBA_2185 | ScienCell Research Laboratories |

| | | |
|---|---|---|
| | HVMSC t Human Vertebral Mesenchymal Stem Cell, MBA_2187 | ScienCell Research Laboratories |
| | HMF t Human Mammary Fibroblast, MBA_2189 | ScienCell Research Laboratories |
| | HUVSMC t Human Umbilical Vein Smooth Muscle Cell, MBA_2192 | ScienCell Research Laboratories |
| | HUASMC t Human Umbillical Artery Smooth Muscle Cell, MBA_2193 | ScienCell Research Laboratories |
| | HEM-l t Human Epidermal Melanocyte, MBA_2116 | ScienCell Research Laboratories |
| | HEM-m t Human Epidermal Melanocyte-medium, MBA_2117 | ScienCell Research Laboratories |
| | HEM-d t Human Epidermal Melanocyte -dark, MBA_2118 | ScienCell Research Laboratories |
| | DPSC P6 ctrl | Songtau Shi School of Dentistry USC |
| | 1837 HPLF P5 (human peridontal ligament) | ScienCell Research Laboratories |
| | DPSC (Dental Pulp Stem Cells) | Songtau Shi School of Dentistry USC |
| | Bone marrow mesenchymal stem cells (PromoCell) P7 ctrl | PromoCell |
| | Human brain vascular smooth muscle cells (SciceCell lot#4762) P7 ctrl | ScienCell Research Laboratories |
| | HAoSMC (Human aortic smooth muscle cells) | PromoCell |
| | HAoSMC (aortic smooth muscle PromoCell) lot 3992005 P7 ctrl | PromoCell |
| | HBMEC tRNA (Human Brain Microvascular Endothelial Cell total RNA), 10 µg | ScienCell Research Laboratories |
| | HCPEC tRNA (Human Choroid Plexus Endothelial Cell total RNA), 10 µg | ScienCell Research Laboratories |
| | HDMEC tRNA (Human Dermal Microvascular Endothelial Cell total RNA),10 µg | ScienCell Research Laboratories |
| | HDLEC tRNA (Human Dermal Lymphatic Endothelial Cell total RNA), 10 µg | ScienCell Research Laboratories |
| | HLEC tRNA (Human Lymphatic Endothelial Cell total RNA), 10 µg | ScienCell Research Laboratories |
| | HIMEC tRNA (Human Intestinal Microvascular Endothelial Cell total RNA), 10 µg | ScienCell Research Laboratories |
| | HPMEC tRNA (Human Pulmonary Microvascular Endothelial Cell total RNA), 10 µg | ScienCell Research Laboratories |
| | HPAEC tRNA (Human Pulmonary Artery Endothelial Cell total RNA), 10 µg | ScienCell Research Laboratories |
| | HRGEC tRNA (Human Renal Glomerular Endothelial Cell total RNA), 10 µg | ScienCell Research Laboratories |
| | HHSEC tRNA (Human Hepatic Sinusoidal Endothelial Cell total RNA), 10 µg | ScienCell Research Laboratories |
| | HCMEC tRNA (Human Cardiac Microvascular Endothelial Cell total RNA), 10 µg | ScienCell Research Laboratories |
| | HAEC tRNA (Human Aortic Endothelial Cell total RNA), 10 µg | ScienCell Research Laboratories |
| | HOMEC tRNA (Human Ovarian Microvascular Endothelial Cell total RNA), 10 µg | ScienCell Research Laboratories |
| | HUVEC tRNA (Human Umbilical Vein Endothelial Cell total RNA), 10 µg | ScienCell Research Laboratories |
| | 1457 HUVEC P6 | Lonza |
| | HUAEC (Human Umbilical Artery Endothelial Cell total RNA), 10 µg | ScienCell Research Laboratories |

| | | |
|---|---|---|
| | HAEC (Lonza human aortic endothelial cells) P6 ctrl | Lonza |
| | HBdMEC t Human Bladder Microvascular Endothelial Cell, MBA_2150 | ScienCell Research Laboratories |
| | HMVEC (Human mammary vascular endothelial cells) P6 | Lonza |
| | Normal human hepatocyte P 35yr old (Celsis In vitro Technologies Product#M0099 ) | Celsis In Vitro Technologies |
| | Normal human hepatocyte P5 64 yr old (Zenbio-HPNP2 lot ZBH2134) | Zenbio |
| | Normal human hepatocyte P5 64 yr old (Zenbio-HPNP2 lot ZBH2134) | Zenbio |
| | HH t Human Hepatocyte | ScienCell Research Laboratories |
| **Neuronal cell Types (NCs)** | HN t Human Neuron | ScienCell Research Laboratories |
| | NHA (normal human astrocytes Lonza) P3 ctrl | ScienCell Research Laboratories |
| | HA t Human Astrocyte | ScienCell Research Laboratories |
| | HA-c t Human Astrocyte-cerebellar | ScienCell Research Laboratories |
| | HA-h t Human Astrocyte-hippocampal | ScienCell Research Laboratories |
| **Diverse human Adult Epithelial Cell types (AECs)** | ATCC MCF 10A Fibrocystic disease (epithelial) | American Type Culture Collection (ATCC) |
| | Human Oral Keritinocytes HOK P5 | ScienCell Research Laboratories |
| | HOK t Human Oral Keratinocyte, MBA_2126 | ScienCell Research Laboratories |
| | HEEpiC t Human Esophageal Epithelial Cell, MBA_2128 | ScienCell Research Laboratories |
| | HPAEpiC t Human Pulmonary Alveolar Epithelial Cell, MBA_2136 | ScienCell Research Laboratories |
| | HBEpiC t Human Bronchial Epithelial Cell, MBA_2137 | ScienCell Research Laboratories |
| | HTEpiC t Human Tracheal Epithelial Cell, MBA_2138 | ScienCell Research Laboratories |
| | HSAEpiC tHuman Small Airway Epithelial Cell, MBA_2139 | ScienCell Research Laboratories |
| | HCPEpiC t Human Choroid Plexus Epithelial Cell, MBA_2104 | ScienCell Research Laboratories |
| | HRPTEpiC t Human Renal Proximal Tubular Epithelial Cell, MBA_2147 | ScienCell Research Laboratories |
| | HREpiC t Human Renal Epithelial Cell, MBA_2148 | ScienCell Research Laboratories |
| | HUC t Human Urothelial Cell, MBA_2152 | ScienCell Research Laboratories |
| | HMEpiC t Human Mammary Epithelial Cell, MBA_2188 | ScienCell Research Laboratories |
| | HPrEpiC r Human Prostate Epithelial Cells, MBA_2153 | ScienCell Research Laboratories |
| | NHEK-neo ctrl P3 | ScienCell Research Laboratories |
| | HEK t Human Epidermal Keratinocyte, MBA_2114 | ScienCell Research Laboratories |
| | HEK-a t Human Epidermal Keratinocyte-adult, MBA_2115 | ScienCell Research Laboratories |
| | NHBE P4 ctrl | ScienCell Research Laboratories |
| | HCEpiC t Human Corneal Epithelial Cell, MBA_2167 | ScienCell Research Laboratories |
| | HK t Human Keratocyte, MBA_2168 | ScienCell Research Laboratories |
| | HRPEpiC t Human Retinal Pigment Epithelial Cell, MBA_2169 | ScienCell Research Laboratories |
| | HLEpiC t Human Lens Epithelial Cell, MBA_2170 | ScienCell Research Laboratories |

| | | |
|---|---|---|
| | HIPEpiC t Human Iris Pigment Epithelial Cell t, MBA_2171 | ScienCell Research Laboratories |
| | HNPCEpiC t Human Non-Pigment Ciliary Epithelial Cell, MBA_2173 | ScienCell Research Laboratories |
| **Diverse sarcoma cell lines** | ATCC TE 159 T rhabdomyosarcoma | American Type Culture Collection (ATCC) |
| | ATCC A 204 rhabdomyosarcoma muscle | American Type Culture Collection (ATCC) |
| | ATCC TE 617 T rhabdomyosarcoma connective tissue | American Type Culture Collection (ATCC) |
| | ATCC Hs 729T rhabdomyosarcoma | American Type Culture Collection (ATCC) |
| | ATCC metastatic rhabdomyosarcoma | American Type Culture Collection (ATCC) |
| | ATCC TE-925.T rhabdomyosarcoma | American Type Culture Collection (ATCC) |
| | ATCC SJCRH30 rhabdomyosarcoma | American Type Culture Collection (ATCC) |
| | ATCC Hs 926.T Renal rhabdomyosarcoma | American Type Culture Collection (ATCC) |
| | ATCC Hs 94T rhabdomyosarcoma | American Type Culture Collection (ATCC) |
| | ATCC TE381.T rhabdomyosarcoma | American Type Culture Collection (ATCC) |
| | ATCC SK-LMS-1 leiomysarcoma/vulva | American Type Culture Collection (ATCC) |
| | ATCC A-673 Ewings sarcoma (muscle) | American Type Culture Collection (ATCC) |
| | ATCC Hs 822.T Ewing's sarcoma | American Type Culture Collection (ATCC) |
| | ATCC Hs 863.T Ewing's sarcoma | American Type Culture Collection (ATCC) |
| | ATCC RD-ES Ewing's sarcoma (bone) | American Type Culture Collection (ATCC) |
| | ATCC G401 rhabdoid sarcoma (derived from Wilms tumor) | American Type Culture Collection (ATCC) |
| | ATCC HOS osteosarcoma | American Type Culture Collection (ATCC) |
| | ATCC KHOS 240S osteosarcoma | American Type Culture Collection (ATCC) |
| | ATCC KHOS NP R 970 S osteosarcoma | American Type Culture Collection (ATCC) |
| | 143B osteosarcoma | American Type Culture Collection (ATCC) |
| | MNNG/HOS Cl #5 R-1059-D osteosarcoma | American Type Culture Collection (ATCC) |
| | MG-63 osteosarcoma | American Type Culture Collection (ATCC) |
| | ATCC U 2 OS osteosarcoma bone ALT | American Type Culture Collection (ATCC) |
| | Saos-2 osteocarcinoma | American Type Culture Collection (ATCC) |
| | ATCC SJSA 1 osteosarcoma multipotential sarcoma bone | American Type Culture Collection (ATCC) |
| | ATCC Hs 737.T giant cell sarcoma (disease: bone) | American Type Culture Collection (ATCC) |
| | ATCC Hs 706.T bone giant cell sarcoma | American Type Culture Collection (ATCC) |
| | ATCC Hs 821.T giant cell sarcoma (bone) | American Type Culture Collection (ATCC) |
| | SW 1353 [SW 1353, SW-1353] chondrosarcoma | American Type Culture Collection (ATCC) |
| | SW-982 synovial sarcoma | American Type Culture Collection (ATCC) |
| | 94T778 liposarcoma | American Type Culture Collection (ATCC) |
| | 93T449 liposarcoma | American Type Culture Collection (ATCC) |
| | SW-872 liposarcoma | American Type Culture Collection (ATCC) |
| | HT-1080 fibrosarcoma | American Type Culture Collection (ATCC) |
| | ATCC MES-SA uterine sarcoma | American Type Culture Collection (ATCC) |
| | ATCC MES-SA uterine sarcoma | American Type Culture Collection (ATCC) |
| | ATCC Hs925.T pagetoid Sarcoma (skin) | American Type Culture Collection (ATCC) |
| | ATCC Hs132T spindle cell sarcoma | American Type Culture Collection (ATCC) |
| | ATCC VA-ES-BJ epithelial sarcoma | American Type Culture Collection (ATCC) |
| | ATCC SK-HEP-1 adenocarcinoma (endothelial) | American Type Culture Collection (ATCC) |
| | ATCC NCI N87 | American Type Culture Collection (ATCC) |
| | ATCC ACHN renal cell adenocarcinoma | American Type Culture Collection (ATCC) |
| | ATCC DLD-1 colorectal adenocarcinoma | American Type Culture Collection (ATCC) |
| | ATCC SK-PN-DW Retroperitoneal Embryonal Tumor Disease: Malignant Primitive Neuroectodermal Tumor | American Type Culture Collection (ATCC) |

| | | |
|---|---|---|
| Diverse carcinoma/adehnocarcinoma cell lines | ATCC DU 145 carcinoma (epithelial morphology) | American Type Culture Collection (ATCC) |
| | ATCC SK-LU-1 adenocarcinoma | American Type Culture Collection (ATCC) |
| | ATCC SCC-4 tongue epithelial-like keritinocyte squamous cell carcinoma | American Type Culture Collection (ATCC) |
| | ATCC PC-3 metastatic adenocarcinoma | American Type Culture Collection (ATCC) |
| | ATCC BT-20 mammary gland carcinoma | American Type Culture Collection (ATCC) |
| | ATCC LNCaP carcinoma (epithelial) | American Type Culture Collection (ATCC) |
| | ATCC M059J Malignant Glioblastoma; Glioma | American Type Culture Collection (ATCC) |
| | ATCC AU565 Adenocarcinoma | American Type Culture Collection (ATCC) |
| | ATCC T-47D ductal carcinoma (epithelial | American Type Culture Collection (ATCC) |
| | ATCC MCF7 Adenocarcinoma (eptithelial) | American Type Culture Collection (ATCC) |
| | ATCC SK-BR-3 adenocarcinoma | American Type Culture Collection (ATCC) |
| | ATCC A-431 epidermoid carcinoma | American Type Culture Collection (ATCC) |
| | ATCC NCI-H358 bronchioalveolar carcinoma; non-small cell lung cancer | American Type Culture Collection (ATCC) |
| | ATCC NCI-H2126 Adenocarcinoma; Non-Small Cell Lung Cancer (epithelial) | American Type Culture Collection (ATCC) |
| | ATCC KLE uterus endometrium adenocarcinoma | American Type Culture Collection (ATCC) |
| | ATCC WiDr Colorectal Adenocarcinoma (epithelial) | American Type Culture Collection (ATCC) |
| | ATCC PA-1 Teratocarcinoma | American Type Culture Collection (ATCC) |
| | ATCC SK-N-DZ Neuroblastoma | American Type Culture Collection (ATCC) |
| | ATCC LS513 Dukes' Type C,Colorectal Carcinoma (epithelial) | American Type Culture Collection (ATCC) |
| | ATCC RKO carcinoma (epithelial) | American Type Culture Collection (ATCC) |
| | ZR-75-1 ductal carcinoma (epithelial) | American Type Culture Collection (ATCC) |
| | MDA-MB-231 Adenocarcinoma (epithelial) | American Type Culture Collection (ATCC) |
| | THLE-3 epithelial cells (liver) transformed with SV40 large T antigen (nontumorigenic) | American Type Culture Collection (ATCC) |
| | ATCC Hep G2 Hepatocellular Carcinoma | American Type Culture Collection (ATCC) |
| | ATCC SNU-387 Grade Iv/V, Pleomorphic Hepatocellular Carcinoma | American Type Culture Collection (ATCC) |
| | ATCC HEB 3B2.1-7 Hepatocellular Carcinoma | American Type Culture Collection (ATCC) |
| | HMepiC human mammary epithelial cell ScienCell | ScienCell Research Laboratories |
| | HTB-133 breast epithelial adenocarcinoma (ATCC) | American Type Culture Collection (ATCC) |
| | HPrEpiC human prostate epithelial line ScienCell | ScienCell Research Laboratories |
| | CRL-1740 LNCaP human prostate epithelial adenocarcinoma (ATCC) | American Type Culture Collection (ATCC) |
| Blood Cell Cancer Lines | ATCC U-937 histiocytic lymphoma | American Type Culture Collection (ATCC) |
| | ATCC JM-1 B-cell lymphoblast lymphoma | American Type Culture Collection (ATCC) |
| | ATCC Kasumi-8 B-cell lymphoblastic leukemia | American Type Culture Collection (ATCC) |
| | HL-60 promyelocytic leukemia | American Type Culture Collection (ATCC) |
